# Somatic regulation of female germ cell regeneration and development in planarians

**DOI:** 10.1101/2021.11.22.469583

**Authors:** Umair W. Khan, Phillip A. Newmark

## Abstract

Female germ cells develop into oocytes, with the capacity for totipotency. In most animals, these remarkable cells are specified during development and cannot be regenerated. By contrast, planarians, known for their regenerative prowess, can regenerate germ cells. To uncover mechanisms required for female germ cell development and regeneration, we generated gonad-specific transcriptomes and identified genes whose expression defines progressive stages of female germ cell development. Strikingly, early female germ cells share molecular signatures with the pluripotent stem cells driving planarian regeneration. We uncovered spatial heterogeneity within somatic ovarian cells and found that a regionally enriched *FoxL* homolog is required for oocyte differentiation, but not specification, suggestive of functionally distinct somatic compartments. Unexpectedly, a neurotransmitter-biosynthetic enzyme, AADC, is also expressed in somatic gonadal cells, and plays opposing roles in female and male germ cell development. Thus, somatic gonadal cells deploy conserved factors to regulate germ cell development and regeneration in planarians.

## Introduction

Germ cells serve as a link between generations, producing the gametes that propagate the genetic material from parent to offspring. During their development, germ cells undergo dramatic, sex-specific differentiation to generate highly specialized cell types, the sperm and egg, which will ultimately yield a totipotent zygote. As the female germline differentiates into oocytes, it acquires the capacity for totipotency (Reik and Surani, 2015; Seydoux and Braun, 2006). This capacity is exemplified by somatic cell nuclear transfer, in which oocyte cytoplasm reprograms differentiated cell nuclei to produce viable clones (Campbell et al., 1996; Gurdon, 1962; Wakayama et al., 1998). Parthenogenesis, in which an unfertilized egg generates an entire new organism (Simon et al., 2003), provides another striking example of the oocyte’s capacity for totipotency. Given the oocyte’s critical roles in embryonic development, understanding the mechanisms underlying female germ cell development has enormous significance.

Somatic support cells within the gonads play critical roles in regulating germ cell development across the animal kingdom. Soma-germline communication is necessary throughout a germ cell’s life, regulating fate choices and survival, to proliferation and differentiation (Kiger et al., 2000; Kimble and White, 1981; Korta and Hubbard, 2010; Li and Albertini, 2013; Murray et al., 2010). The in vitro derivation of functional oocytes and spermatid-like cells from pluripotent stem cells requires co-culture with somatic gonadal cells (Hikabe et al., 2016; Zhou et al., 2016), further emphasizing the importance of soma-germ cell interactions in facilitating proper germ cell development.

Most animals, including model organisms typically used to study animal development, set aside germ cells early during embryonic development and cannot replace them if lost (Nieuwkoop and Sutasurya, 1979, 1981). By contrast, freshwater planarians demonstrate the striking ability to regenerate both male and female germ cells from tissues lacking any reproductive structures (Morgan, 1901; Sato et al., 2006; Wang et al., 2007). Planarians are best known for their remarkable whole-body regeneration, driven by stem cells called neoblasts (Baguñà et al., 1989), a subset of which are pluripotent (Wagner et al., 2011; Zeng et al., 2018). Planarian germ cells are specified inductively and are derived from neoblasts (Baguñà et al., 1989; Newmark et al., 2008; Sato et al., 2006; Wang et al., 2007). The vast majority of research on planarian regeneration focuses on asexual strains that reproduce by fission; however, planarians can also reproduce sexually as simultaneous hermaphrodites. Germ cell development in sexual planarians is responsive to physiological cues: following prolonged starvation, gonads regress and accessory reproductive organs are resorbed (Berninger, 1911; Morgan, 1901; Schultz, 1904); when feeding is resumed, gonads again produce gametes and accessory reproductive organs are re-formed. The remarkable developmental plasticity of the planarian reproductive system provides a unique opportunity to investigate mechanisms regulating sex-specific germ cell specification and differentiation in the context of an adult organism.

In sexual planarians, ovarian tissue is scarce relative to testes; as such, previous transcriptomic analyses of the planarian reproductive system predominantly identified genes enriched in the testes rather than ovaries (Rouhana et al., 2017; Wang et al., 2010; Zayas et al., 2005). Whole-animal single-cell sequencing approaches produced similar results (Fincher et al., 2018). Thus, little is known about the gene-expression changes driving female germ cell development in planarians. An orphan G-protein-coupled receptor (GPCR), Ophis, is expressed in somatic gonadal cells of ovaries and testes and is required for differentiation of both oocytes and sperm (Saberi et al., 2016); however, gene-expression differences distinguishing female from male somatic gonadal cells have yet to be identified, and the roles of somatic ovarian cells in various stages of female germ cell development remain to be determined.

To circumvent the limited quantity of ovarian tissue and identify genes with enriched expression in planarian ovaries, we generated gonad-specific transcriptomes using laser-capture microdissection followed by RNA sequencing (RNA-seq). Analysis of the resulting transcriptomes and validation by in situ hybridization identified genes defining progressive stages of female germ cell development and revealed similarities between female germ cells and pluripotent neoblasts. These studies also discovered spatial heterogeneity in the ovarian somatic cell population; RNA interference (RNAi) targeting a planarian homolog of *FoxL2*, a regulator of granulosa cell identity in mammalian ovaries, suggests that this heterogeneity reflects distinct functional somatic compartments within the ovary. Finally, we found that a gene encoding a monoamine-neurotransmitter-synthetic enzyme is expressed in somatic gonadal cells and plays opposing roles in male and female germ cell development, suggesting non-neuronal roles for monoamine transmitters in regulating germ cell development and regeneration in planarians.

## Results

### LCM-Seq identifies genes that mark and distinguish ovaries from testes

The reproductive system of the sexual strain of the planarian *Schmidtea mediterranea* consists of a pair of ovaries located ventrally under the posterior lobes of the brain, numerous dorsolaterally distributed testes, and accessory reproductive organs (yolk glands (vitellaria), oviducts, sperm ducts, copulatory apparatus, and gonopore) (Fig. 1A) (Hyman, 1951; Issigonis and Newmark, 2019). To precisely excise ovarian, testis, and surrounding non-gonadal tissues for transcriptomic comparisons, we used laser-capture microdissection (LCM; Fig. 1B)(Emmert-Buck et al., 1996; Espina et al., 2006; Forsthoefel et al., 2020). RNA-seq analysis (Mortazavi et al., 2008; Nagalakshmi et al., 2008) of the laser-captured tissues identified 36,036 non-redundant transcripts, of which 7,557 (21%) were upregulated (≥2-fold, FDR p-value ≤ 0.01) in either or both gonads: 1,491 (4%) in ovaries; 4,880 (14%) in testes; and 1,206 (3%) in both (Fig. 1B-C, Fig S1A-D, Table S1). To validate the LCM-generated gonadal transcriptomes, we examined genes with previously reported gonadal expression (Wang et al., 2010). As expected, we found significant upregulation of the majority of these genes (82/89) in the testis transcriptome (Fig. S2A) including *nanos,* a conserved regulator of germ cell development (Handberg-Thorsager and Saló, 2007; Sato et al., 2006; Wang et al., 2007), which was also upregulated in the ovary transcriptome (Fig. S2A).

**Figure 1.**
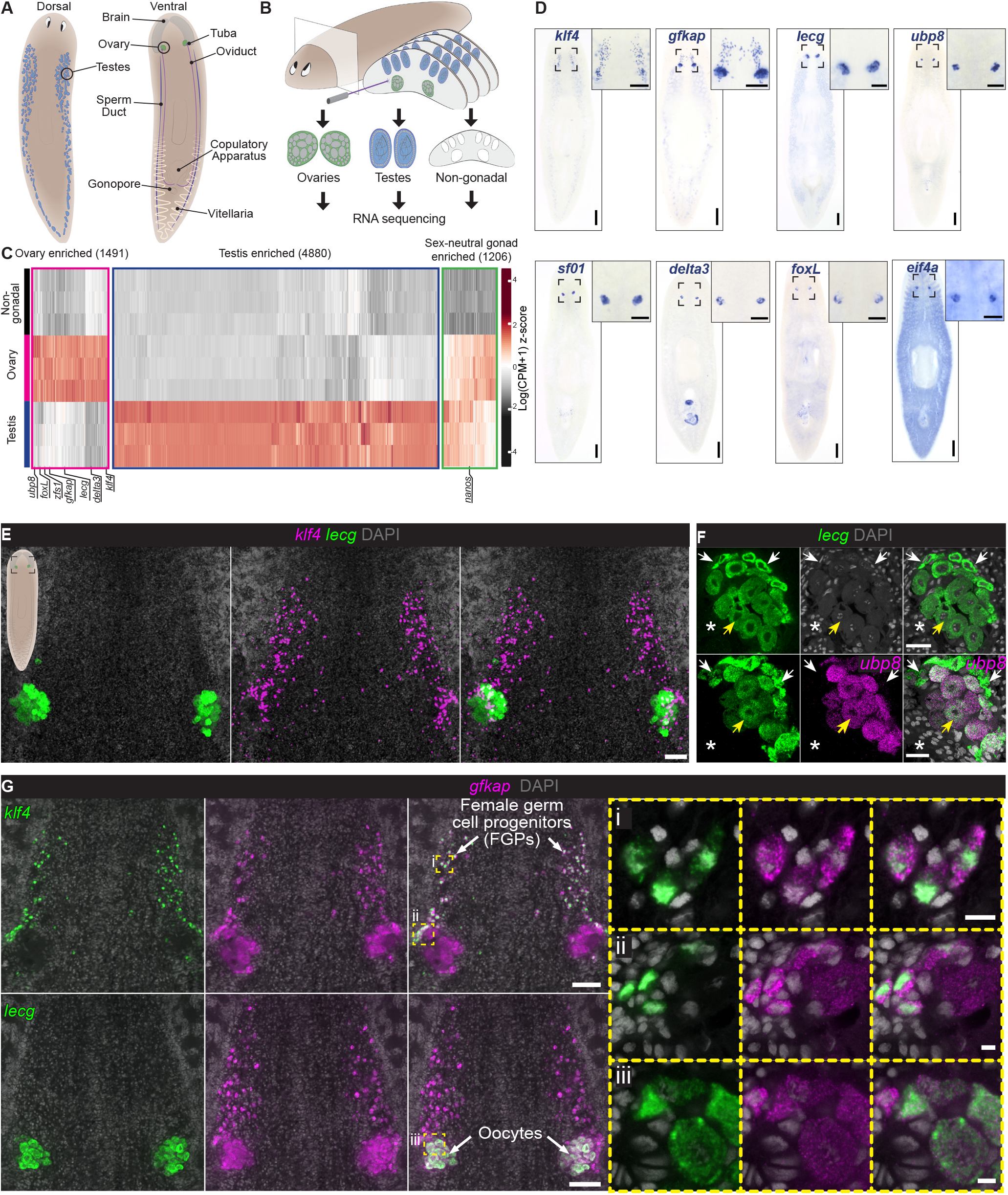
Ovary-specific transcriptome reveals progressive stages of female germ cell development. (A) Reproductive system of the sexual strain of *S. mediterranea*. (B) LCM-RNAseq approach to generate gonadal transcriptomes. (C) Hierarchical clustering of genes with significantly enriched expression in ovaries, testes, or both. (D) Representative whole-mount RNA in situ hybridizations (WISH) to detect candidate ovary-enriched transcripts. (E) Double fluorescent RNA in situ hybridization (FISH) to detect *klf4* and *lecg* reveals a field of *klf4^+^* cells anterior to the ovary. (F) FISH to detect *lecg* and *ubp8*, revealing expression in oocytes. Smaller immature oocytes (white arrows) are observed at the periphery of the ovary, while larger mature oocytes (yellow arrows) reside internally, proximal to the tuba (fertilization duct) (asterisks). (G) FISH to detect *gfkap* and *klf4* or *lecg* shows *gfkap* expression in *klf4^+^* cells anterior to the ovary (i) and at the margin of the ovary (ii). Co-expression of *gfkap* with *lecg* is observed in oocytes (iii). *klf4^+^gfkap^+^* cells are presumptive female germ cell progenitors (FGPs) that differentiate into *gfkap^+^lecg^+^* oocytes within the ovary. Nuclei are labeled with DAPI. Scale bars: (D) 500 μm; insets: 200 μm; (E) 100 μm; (F) 50 μm; (G) 100 μm; insets: 10 μm. See also Figures S1 and S2.

To validate our list and identify new genes expressed in the ovary, we performed whole-mount colorimetric RNA in situ hybridization (WISH) for ovary-enriched transcripts and detected diverse expression patterns in and around the ovaries (Fig. 1D, Fig. S2B). Approximately 90% (204/227) of genes tested showed enriched expression in the ovary or associated reproductive structures. These results demonstrate the successful application of LCM-RNA-seq for generating gonad-specific transcriptomes, providing a resource for identifying regulators of germ cell development and regeneration.

### Female germ cell progenitors (FGPs) are specified outside the ovaries and express markers of pluripotency

Histological and ultrastructural studies have shown that planarian oocytes grow and mature in a progression from the margins of the ovary, toward the tuba (fertilization duct) (Gremigni and Nigro, 1983; Harrath et al., 2011). Markers of the earliest stages of germ cell development (*nanos* and *krüppel-like factor 4* (*klf4*)) label oogonia at the ovary periphery (Issigonis et al., 2021; Wang et al., 2007); *nanos* and *klf4* are also expressed in two fields of cells, anterior to the ovary at the ventro-medial portion of each brain lobe (Issigonis et al., 2021). To help elucidate the origins and differentiation of the female germ cell lineage, we sought to identify ovary-enriched transcripts expressed at various stages of oogenesis. Fluorescent RNA in situ hybridization (FISH) analysis detected *galactose-binding lectin* (*lecg*) expression in differentiating cells throughout the ovary, but not in the anterior field of *klf4*-expressing cells (Fig. 1E). Expression of *ubiquitin carboxyl-terminal hydrolase 8* (*ubp8*) was detected in larger, more mature oocytes residing within the ovary, proximal to the tuba, but not within smaller *lecg*+ cells at the ovary periphery (Fig. 1F). Thus, *ubp8* is upregulated later in oocyte differentiation than *lecg* (Fig. 1F). The expression patterns of these oocyte markers and the corresponding growth of oocytes reveal progressive stages of female germ cell differentiation, from the periphery of the ovary, towards its interior.

Like the previously described patterns of *nanos* and a homolog of the pluripotency factor *klf4* (Issigonis et al., 2021), we detected expression of a gene originally annotated as *polyketide synthase-1* (*pks1*) (Zeng et al., 2018) in fields of cells anterior to the ovary; its expression was also detected throughout the ovary (Fig. 1D, 1G). This gene was identified via single-cell sequencing as a marker of a pluripotent subpopulation of neoblasts in asexual *S. mediterranea* (Zeng et al., 2018); because its predicted product shares no conserved domains with any reported proteins, including polyketide synthases, we propose renaming it as the *gene formerly known as pks1 (gfkap)*. *Gfkap* is expressed in the *klf4^+^* fields anterior to the ovaries (Fig. 1G, 1G(i)) and around the margins of the ovary (Fig. 1G, 1G(ii)). The *gfkap*^+^ cells in the anterior fields and around the margins of the ovary were labeled by anti-phospho-Histone H3 (pHH3) antibodies (Fig. S3A(i) and (ii)), indicating that they are proliferative. *Gfkap* is also expressed in *lecg^+^* oocytes within the ovary (Fig. 1G, 1G(iii)), and thus, serves as a marker spanning all stages of female germ cell development. These results suggest that the *nanos^+^ klf4^+^ gfkap^+^* cells located at the margin of the ovary are oogonia that give rise to *gfkap^+^lecg^+^* oocytes within the ovary (Fig. 1G). Finding proliferating cells expressing the earliest markers of germ cell development (*nanos^+^ klf4^+^gfkap^+^*) anterior to the ovaries suggests that female germ cell progenitors (FGPs) can also be specified outside of the ovaries (Issigonis et al., 2021). Identifying markers for progressive stages of female germ cell development allows us to study early steps in female germ cell specification and differentiation.

### A marker of pluripotent neoblasts is expressed in germ cells and only upregulated in neoblasts after wounding

In addition to *gfkap,* single-cell analysis of a pluripotent neoblast sub-population identified other cluster-defining transcripts, including *tspan group-specific gene-1* (*tgs-1)* and *tetraspanin-1 (tspan-1)* (Zeng et al., 2018). The expression of *gfkap* in FGPs of sexual planarians suggested affinities between pluripotent neoblasts and early germ cells; thus, we sought to further explore this relationship. First, we examined the expression of the neoblast marker *piwi-1* (Reddien et al., 2005), a planarian PIWI homolog that is also expressed in germ cells (Davies et al., 2017). We detected relatively high levels of *piwi-1* mRNA in *gfkap^+^* FGPs (Fig. 2A, 2A(i-ii)). Consistent with previous studies (Davies et al., 2017), we detected *piwi-1* expression in oocytes, suggesting that *piwi-1* expression is maintained through oocyte differentiation (Fig. 2A, 2A(iii)). Similar to *gfkap*, *tgs-1* is enriched in the ovarian transcriptome and expressed abundantly in FGPs (Fig. 2B, Fig. S3C); however, *tgs-1* was also detected in a subset of neoblasts (Fig. S3D). In the sexual strain, *gfkap* expression was most pronounced in germ cells in both ovaries and testes (Fig. 1G, 2A, 2B, Fig. S3B) and, aside from co-expression in the germ cells, its expression did not overlap with the expression of *piwi-1*.

**Figure 2.**
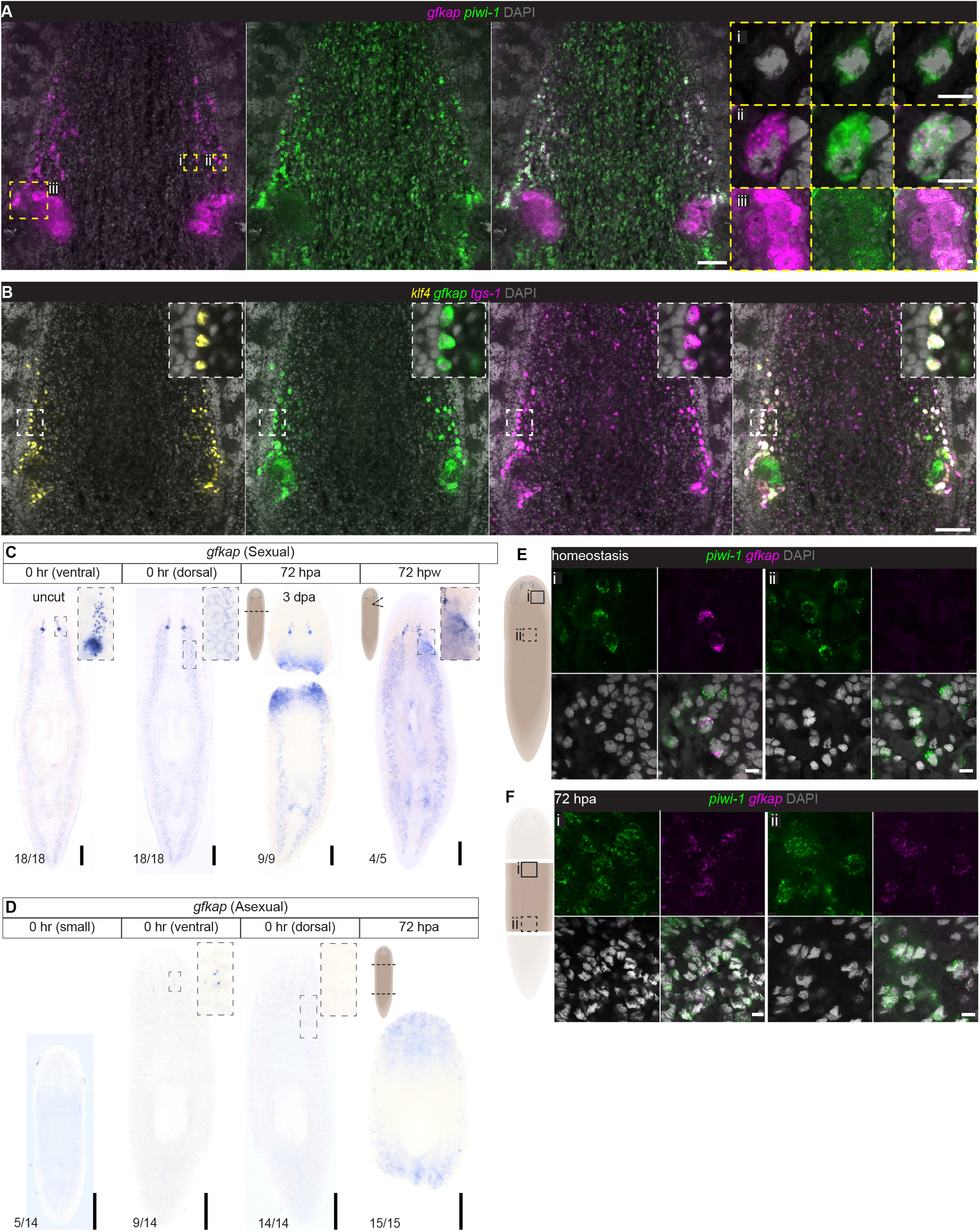
Female germ cell progenitors express markers associated with a pluripotent neoblast subpopulation. (A) FISH to detect *gfkap* and the neoblast marker, *piwi-1.* Expression of *piwi-1* is maintained from neoblasts (i) into FGPs (ii) and oocytes (iii). (B) FISH to detect *tgs-1* with *klf4* and *gfkap*. *Tgs-1* marks a pluripotent subset of neoblasts and is enriched in FGPs. (C, D) WISH to detect *gfkap* mRNA in sexuals (C) and asexuals (D) during homeostasis (uncut) and 72 hours post-amputation (hpa) or -wounding (hpw). *gfkap* is upregulated at wound sites in both sexuals and asexuals. (E, F) FISH to detect *gfkap* and *piwi-1* in asexuals during homeostasis (E) and 72 hpa (F), shows upregulation of *gfkap* in neoblasts at wound sites after amputation. Scale bars: (A-B) 100 μm; insets: 10 μm; (C-D) 500 μm; (E-F) 10 μm.

To validate the expression of neoblast markers in germ cells, we first assessed *gfkap* expression in the asexual strain of *S. mediterranea*. Although these animals reproduce by fission they develop gonad primordia containing germ cells that fail to differentiate (Handberg-Thorsager and Saló, 2007; Wang et al., 2007). In small asexuals (worm length: 2.3±0.2mm) we did not detect *gfkap* expression anywhere in the animal; larger asexuals (4.2±0.5mm), however, displayed a small group of *gfkap^+^* cells ventral to the posterior brain lobes (Fig. 2D and 2E), reminiscent of FGPs in the sexual strain and consistent with the increase of *nanos*^+^ *klf4*^+^ female germ cells in this region of larger asexual planarians (Issigonis et al., 2021). We did not detect *gfkap* expression in testis primordia of asexuals, suggesting that *gfkap* is upregulated during a later stage of male germ cell development (Fig. 2D).

Considering the scarcity of FGPs in asexuals, it seems unlikely that these cells were captured during single-cell sequencing by Zeng et al.; rather, *gfkap* expression in pluripotent neoblasts could result from cell dissociation and the subsequent induction of wound-response transcriptional programs (Lacar et al., 2016; Wu et al., 2017). Indeed, wound responsiveness was one of the criteria used to define this neoblast subcluster (Zeng et al. 2018). To test this idea, we examined *gfkap* expression by WISH after amputating or injuring the worms and found that *gfkap* mRNA was upregulated at the wound sites in both sexual and asexual strains (Fig. 2C-F). We corroborated our ISH analysis by examining previous regeneration transcriptomes (Kao et al., 2013; Rozanski et al., 2019), which revealed that *gfkap* is upregulated as early as 4 to 6 hours after injury: a time frame consistent with the possibility of cell-dissociation-induced upregulation in neoblasts (Fig. S4A). In contrast, other early germ cell markers, such as *klf4* and *nanos*, are not upregulated at wound sites (Fig. S4B-C). Next, we confirmed that *gfkap* was upregulated in neoblasts after amputation by performing double FISH with the neoblast marker *piwi1*. In uninjured asexual planarians, *gfkap^+^piwi1^+^* cells were only detected in the FGP region (Fig. 2E). However, after amputation, *gfkap^+^piwi1^+^* cells were detected at both anterior and posterior wound sites (Fig. 2F). These data indicate that in intact planarians, *gfkap* marks germ cells rather than neoblasts. Wound-inducible expression of this gene in neoblasts provides another link between germ cells and the planarian’s pluripotent stem cells.

### Gene expression profiling reveals spatial heterogeneity in ovarian somatic cells

How are FGPs maintained and what regulates their differentiation within the ovary? Somatic gonadal cells play critical roles in regulating germ cell development throughout the animal kingdom. To date, studies of soma-germline interactions in the planarian ovary have been limited by the availability of appropriate markers. The first such marker, *ophis*, encodes an orphan GPCR expressed in somatic support cells of ovaries and testes and is required for both female and male germ cell differentiation (Saberi et al., 2016). Our ISH screen of ovary-enriched transcripts identified three genes with expression patterns distinct from those observed for germ cell-enriched transcripts, and resembling a scaffold surrounding the germ cells: *delta3*; the forkhead-family transcription factor, *foxL*; and *endothelin converting enzyme 1* (*ece1*) (Fig. 1C and D, Fig. S5A). We confirmed that *delta3, foxL,* and *ece1* transcripts were not detected in female germ cells using double FISH with pooled probes for the germ cell markers *klf4* and *lecg* (Fig. 3A, 3B, 3C). Additionally, unlike *ophis* (Saberi et al., 2016)*, delta3, foxL,* and *ece1* transcripts were not detectable in the testes, indicating sexually dimorphic expression (Fig. S5A). *delta3^+^* cells are closely associated with oocytes, and co-express *ophis* within the ovary, confirming *delta3* as a marker of ovarian somatic cells (Fig. 3D). Double FISH to detect *foxL* and *delta3* revealed higher expression of *foxL* in *delta3^+^* somatic cells situated proximal to the tuba, compared to *delta3^+^* cells at the periphery (Fig. 3E). In contrast, *ece1* is expressed in the peripheral *delta3^+^* cells (Fig. 3F). Double FISH to detect both *ece1* and *foxL* confirmed the presence of *ece1^high^foxL^low^* cells at the periphery of the ovary (Fig. 3G, 3G(i), 3H) and *ece1^low^ foxL^high^* cells proximal to the tuba (Fig. 3G, 3G(ii), 3H), indicating heterogeneity among ovarian somatic support cells. This heterogeneity could reflect distinct functional roles at different stages of oogenesis.

**Figure 3.**
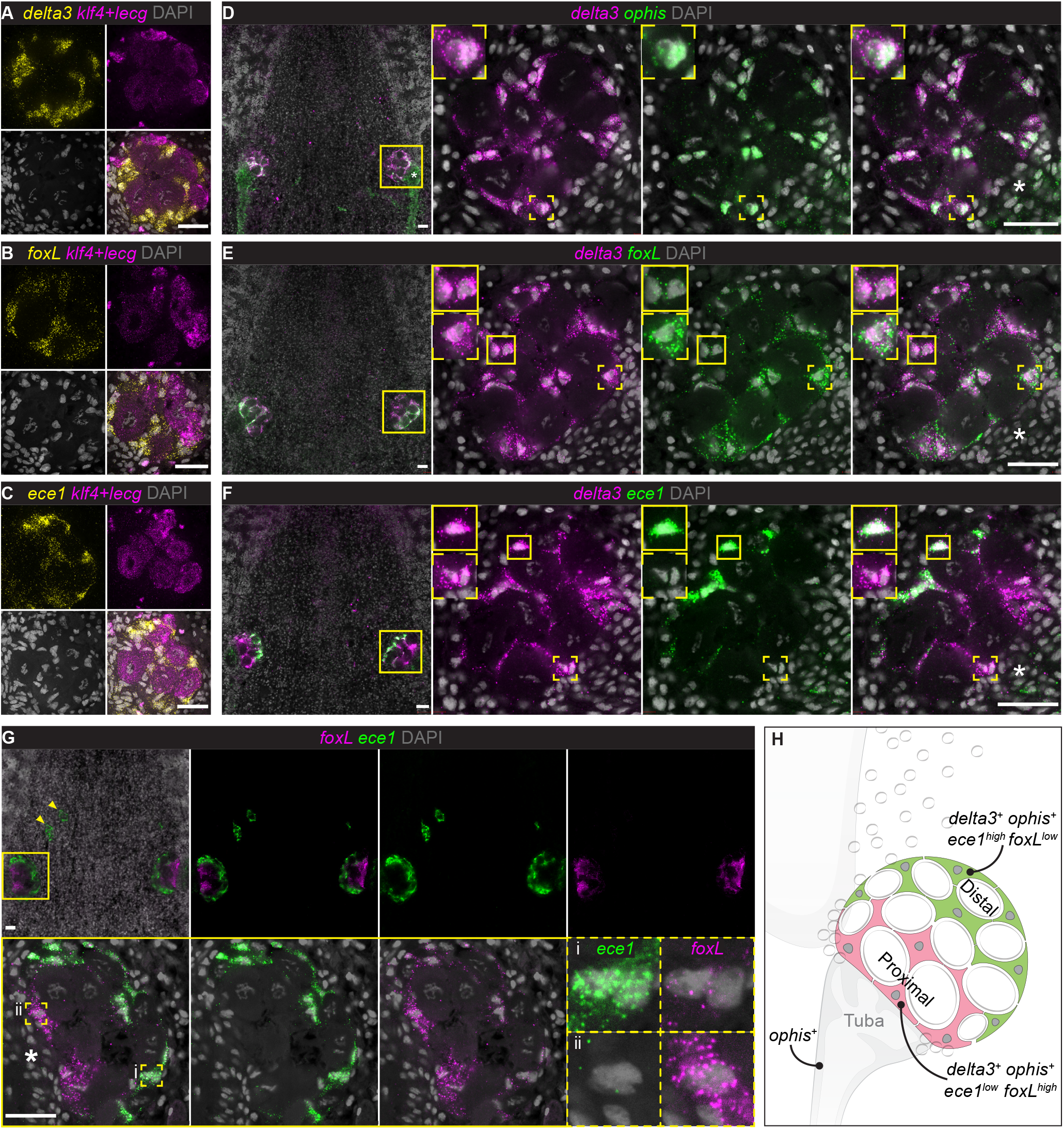
Characterization of somatic ovarian cells. (A, B, and C) FISH to detect *delta3* (A), *foxL* (B), *ece1* (C), and pooled probes for *klf4* (FGPs) and *lecg* (oocytes) shows expression of these transcripts in non-germ cell populations of the ovary. (D) *ophis^+^* ovarian somatic cells co-express *delta3.* (E, F) *delta3^+^* somatic support cells co-express *foxL* (E) and *ece1* (F). (G) FISH to detect *ece1* and *foxL* reveals heterogeneity in gene expression within somatic ovarian cells: (i) *ece1^low^foxL^high^* cells are located proximal to the tuba and (ii) *ece1^high^foxL^low^* cells are located distal to the tuba. Arrowheads indicate ectopic ovaries often found in large sexual worms. (H) Schematic depicting the distribution of and marker gene expression in somatic ovarian cells and associated reproductive structures (tuba and oviduct). Asterisks: tuba. Scale bars: 50 μm. See also Figures S3 and S4.

### Ovarian somatic cells regulate female germ cell development via conserved factors

*FoxL* encodes a planarian homolog of the forkhead-family transcription factor FoxL2 (Fig. S5B), which is a critical regulator of granulosa cell differentiation in the mammalian ovary. Ovaries fail to develop properly in *FoxL2* knockout mice: the absence of functional granulosa cells in these mutants leads to oocyte atresia and infertility due to progressive follicular depletion (Schmidt et al., 2004; Uda et al., 2004). Human mutations in *FoxL2* can lead to premature ovarian failure (POF) or ovarian tumors (Crisponi et al., 2001; Georges et al., 2014; Schmidt et al., 2004; Shah et al., 2009; Uda et al., 2004; Uhlenhaut and Treier, 2011). *FoxL2* is also required for actively maintaining female fate in the adult mouse ovary (Uhlenhaut et al., 2009). Although *FoxL2* genes have been identified in a range of invertebrates, its expression in ovarian somatic cells, but not germ cells, has been proposed to be a vertebrate innovation (Bertho et al., 2016). Furthermore, functional roles for *FoxL2* homologs in ovarian development have yet to be demonstrated in any invertebrate.

Thus, it was noteworthy to find planarian *foxL* similarly expressed in ovarian somatic cells, enriched in the tuba-proximal population surrounding late-stage oocytes (Fig. 3B, 3E, 3G, Fig. S5A). To examine whether *foxL* also plays a functional role in oogenesis, we used RNAi to knock down its expression in the context of decapitated worms that will regenerate new ovaries (Fig. 4A). *foxL(RNAi)* worms regenerated their heads normally and displayed normal feeding behavior (Fig. S5C). The distribution and numbers of *klf4^+^* FGPs in the anterior fields were not affected significantly in these animals (Fig. 4B, 4C). However, *foxL RNAi* resulted in a dramatic and significant reduction in the number of *lecg^+^* oocytes (Fig. 4B, 4C). These data suggest that *foxL* is required cell non-autonomously for oocyte differentiation and maintenance in planarians.

**Figure 4.**
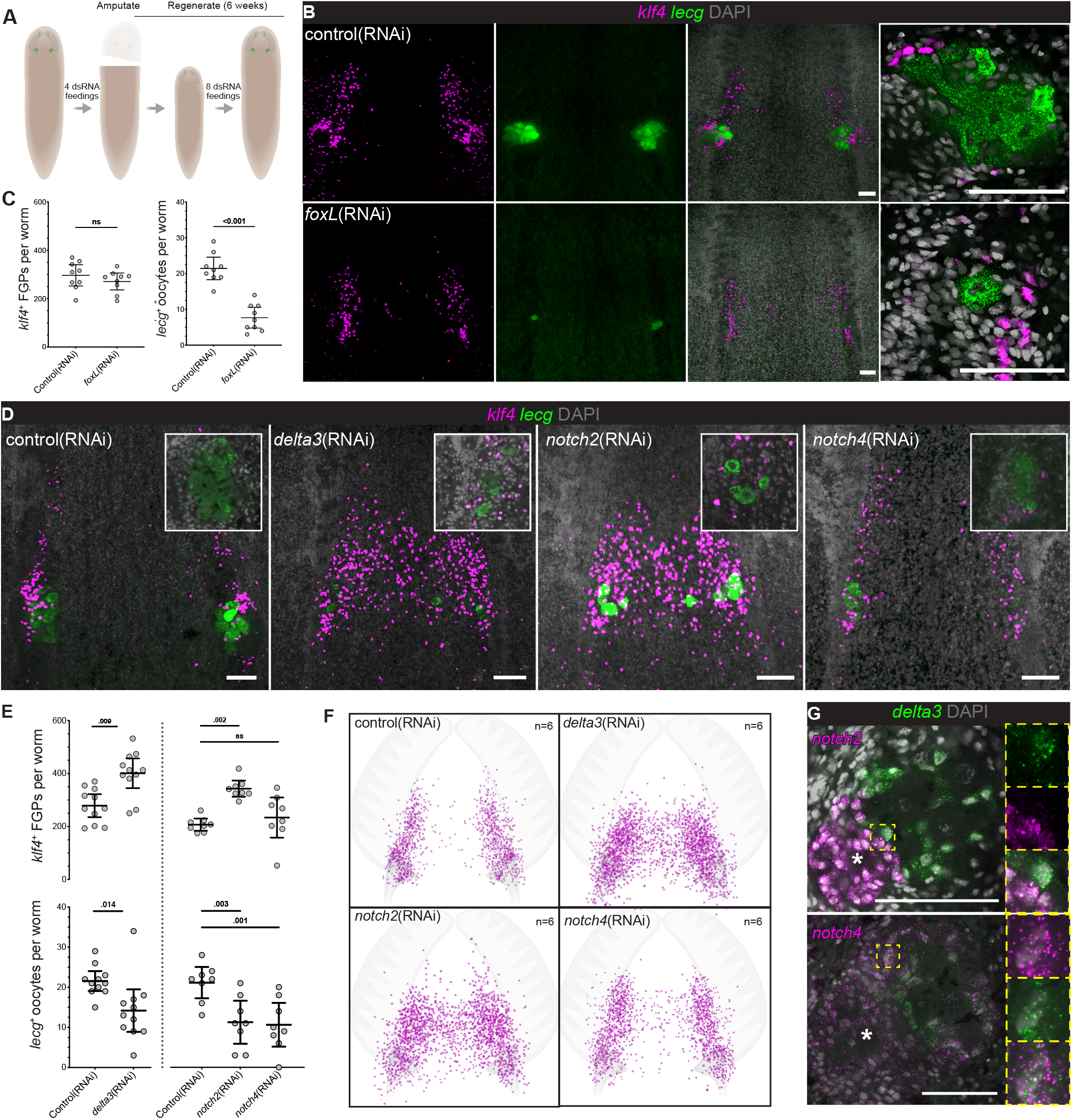
Ovarian somatic cells are essential for female germ cell development. (A) Regeneration assay for RNAi-mediated gene knockdown. (B) FISH to detect *klf4* and *lecg* in control(RNAi) and *foxL*(RNAi) worms. (C) Quantification of *klf4^+^* FGPs and *lecg^+^* oocytes in control(RNAi) and *foxL*(RNAi) worms. (D) FISH to detect *klf4* and *lecg* in control(RNAi), *delta3*(RNAi), *notch2*(RNAi), and *notch4*(RNAi) worms (ns = not significant; mean with 95% CI). (E) Quantification of *klf4^+^* FGPs and *lecg^+^* oocytes in control(RNAi), *delta3*(RNAi), *notch2*(RNAi), and *notch4*(RNAi) worms. (F) Density plot of *klf4*^+^ FGPs in control(RNAi), *delta3*(RNAi), *notch2*(RNAi) and *notch4*(RNAi) worms. The brain lobe and tuba were used to define the relative position of cells. (G) Double FISH for *notch2* or *notch4* with the somatic marker *delta3*. Asterisks: tuba. Scale bars: 100 μm. See also Figure S5.

As reported above, ovarian somatic cells also express a *delta3* homolog (Fig. 3A, 3B), which encodes a transmembrane Notch-signaling ligand (Fig. S5D) (Bray, 2006). Notch signaling plays critical roles in various aspects of soma-germ cell interactions across species. For example, the *C. elegans* niche uses Notch signaling to regulate GSC maintenance and proliferation (Byrd and Kimble, 2009), whereas in the *Drosophila* ovary Notch signaling controls the formation and maintenance of the niche (Xie et al., 2008). To explore the role of Notch signaling in planarian oogenesis, we performed *delta3* RNAi experiments in the context of the ovarian regeneration paradigm (Fig. 4A). *delta3*(*RNAi*) planarians displayed a significant increase of *klf4^+^* FGPs, the distribution of which was skewed toward the midline (Fig. 4D, 4E, 4F). Additionally, the ovaries of *delta3*(*RNAi*) planarians appeared disorganized, with *klf4^+^* FGPs intermingled with *lecg^+^* oocytes; oocyte numbers were significantly reduced (Fig. 4D, 4E).

Because direct cell-cell interaction is necessary for Notch signaling (Bray, 2006), we asked whether Notch receptors were also expressed in the ovary. We found enriched expression of transcripts encoding both planarian Notch receptors in our ovarian transcriptome (Fig. S5D, E). FISH revealed mRNAs of both *notch2* and *notch4* in the tuba and oviduct; *notch4* mRNA was also detected in the *delta3^+^* ovarian somatic cells (Fig. 4G). Knockdown of *notch2* resulted in a significant expansion, midline-skewed distribution of *klf4^+^* FGPs and disorganized ovaries with depleted oocytes, similar to that observed in *delta3(RNAi*) worms. By contrast, *klf4^+^* FGPs appeared unaffected after *notch4* RNAi, but the oocyte numbers were reduced (Fig. 4D, 4E, 4F). We did observe that although *notch2(RNAi)* worms regenerated normally, they were smaller at the end of our assay, suggesting roles in growth or feeding, which could also influence oocyte development (Fig. S5F). Altogether, Notch signaling plays several roles in female germ cell development in planarians, regulating the number and distribution of FGPs, as well as the spatial organization of the ovary, with effects upon oocyte production. The expression patterns of the Notch signaling components reported here suggest that interactions between accessory reproductive organs (oviduct, tuba) and ovarian somatic cells may also help establish proper structure of the ovary.

### A female-specific regulator of germ cell regeneration: bidirectional soma-germline communication

In addition to markers of ovarian somatic cells, the ovary transcriptome enabled us to identify *zfs1,* a female-specific early germ cell marker. This gene encodes a predicted RNA-binding protein (Fig. S6A) and is expressed in ovaries as well as in cells anterior to the ovaries (Fig. 5A, 5B). Double FISH with *gfkap* indicated that *zfs1* is co-expressed in FGPs and oocytes (Fig. 5C); however, unlike other FGP markers (*klf4, nanos,* and *gfkap*) that are also expressed in male germ cells, *zfs1* expression was restricted to female germ cells (Fig. 5A, 5B). RNAi knockdown of *zfs1* using the previously described regeneration paradigm (Fig. 4A) resulted in a dramatic reduction in the number of *klf4^+^* FGPs and ablation of *lecg^+^* oocytes (Fig. 5D, 5E). By contrast, male germ cell development was unaffected by *zfs1* RNAi: *klf4^+^* early male germ cells at the testis periphery and DAPI-stained sperm in the lumen (Issigonis et al., 2021) appeared normal (Fig. 5F). Accessory reproductive structures and egg laying were also unaffected. However, *zfs1(RNAi)* animals were sterile: they failed to produce hatchlings (Fig. S6B-D). These results indicate a germ cell-intrinsic, female-specific role for *zfs1*, which belongs to a small, yet ancient family of RNA-binding proteins that regulate mRNA turnover (Cuthbertson et al., 2008; Wells et al., 2017). The yeast ortholog is a regulator of sexual differentiation (Kanoh et al., 1995; Navarro et al., 2017) and the mammalian homolog, ZFP36L2, is critical for oocyte maturation, targeting transcriptional regulators and chromatin modifiers for degradation (Dumdie et al., 2018). Understanding how *zfs1* expression is restricted to female germ cells will provide insight into the modulation of germ cell sex in a simultaneous hermaphrodite.

**Figure 5.**
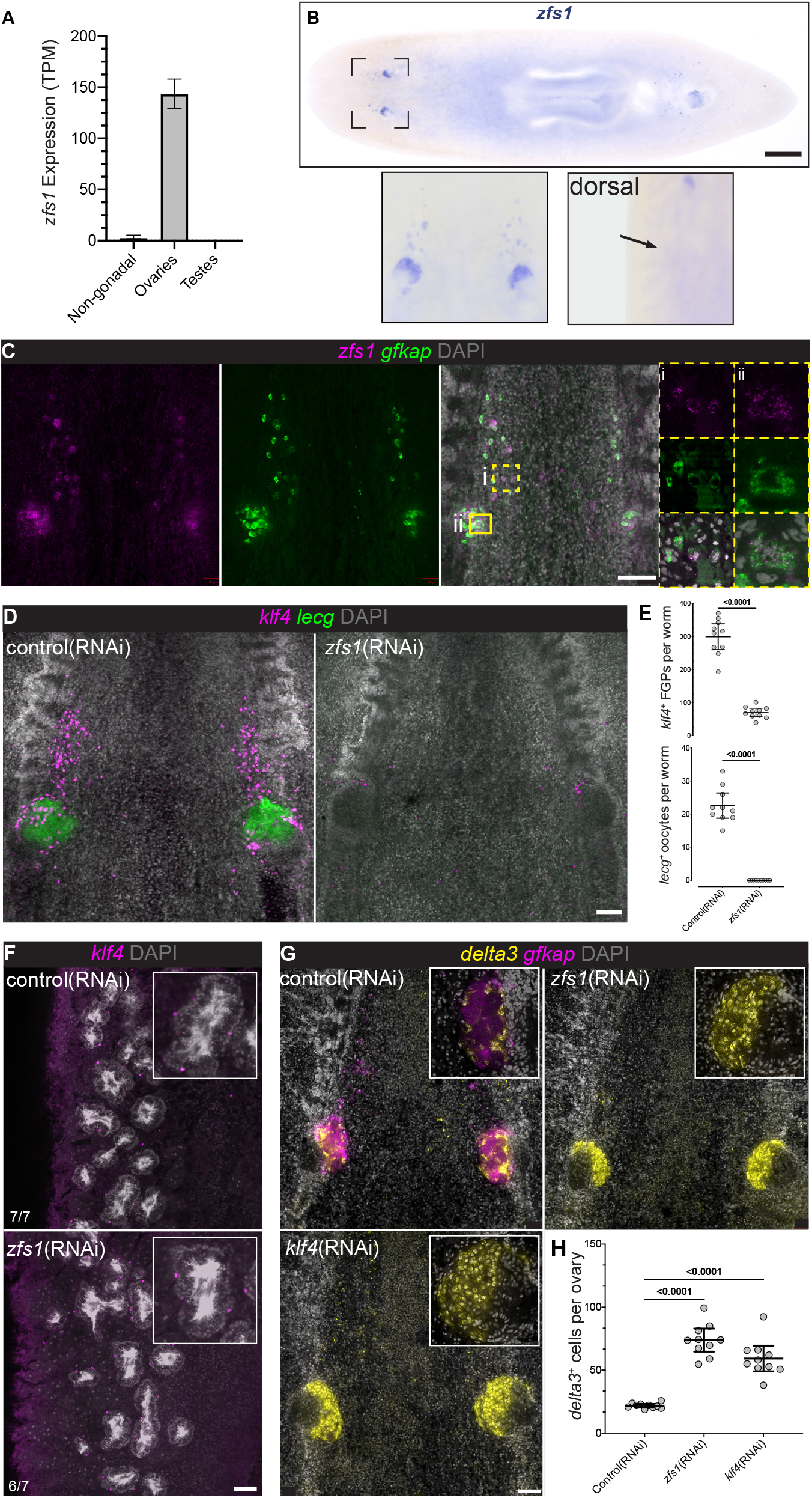
Female-specific role of *zfs1* in germ cell regeneration. (A) *zfs1* expression is enriched in the ovary vs. testis and non-gonadal transcriptomes. (B) WISH to detect *zfs1* reveals expression in the ovary and cells anterior to the ovary. Dorsal view shows no detectable *zfs1* expression in the testes. (C) Double FISH to detect *zfs1* and *gfkap* shows *zfs1* expression in *gfkap*^+^ FGPs anterior to the ovary (i) and oocytes within the ovary (ii). (D) FISH to detect *klf4^+^* FGPs and *lecg^+^* oocytes in control(RNAi) and *zfs1*(RNAi) worms. (E) Quantification of *klf4^+^* FGPs and *lecg^+^* oocytes in control(RNAi) and *zfs1*(RNAi) worms. (F) FISH to detect *klf4* (early male germ cells) in testes of control(RNAi) and *zfs1*(RNAi) worms. In a wild-type testis immature germ cells are located peripherally and differentiating germ cells (spermatids and sperm) are found toward the lumen of each lobe (visible by intense DAPI staining). (G) FISH to detect *delta3* (somatic ovarian cells) and *gfkap* (female germline) in control(RNAi), *zfs1*(RNAi), and *klf4*(RNAi) worms. (H) Quantification of *delta3^+^* support cells in control(RNAi), *zfs1*(RNAi), and *klf4*(RNAi) worms. Scale bars: (B) 500 μm; (C-F) 100 μm. See also Figure S6.

We recently reported that *klf4* RNAi resulted in loss of female germ cells and a concomitant expansion of somatic ovarian cells (Issigonis et al., 2021). Because *klf4* expression is not limited to female germ cells and *klf4* RNAi also affects testis and vitellaria development, this somatic cell expansion could have arisen from altered systemic cues (e.g., due to lack of vitellaria). The ovary-specific effects of *zfs1* RNAi provided a means to test this idea. Using *delta3* as a pan-somatic marker, we observed an increase in somatic support cells in *zfs1*(RNAi) worms similar to that observed after *klf4* RNAi (Fig. 5G, 5H). Thus, the absence of female germ cells triggers an expansion of ovarian somatic cells, suggesting bidirectional communication between germ cells and somatic support cells during ovary regeneration.

### Novel roles of AADC in germ cell development and regeneration

The laser-capture transcriptomes also revealed unanticipated gonadal expression of conserved genes not previously implicated in gonadal function. Aromatic L-amino acid decarboxylase (AADC) is a broadly conserved enzyme (Fig. S7A) that catalyzes a key reaction in the production of monoamine neurotransmitters, such as serotonin and dopamine. Previous studies of asexual planarians showed that *aadc* is expressed in serotonergic neurons within the central nervous system, as well as in pigment cups of the photoreceptors and secretory cells around the pharynx (Currie and Pearson, 2013; März et al., 2013). Thus, it was surprising to find enriched expression of *aadc* mRNA in the planarian ovary (Fig. S7B). We confirmed ovary-enriched *aadc* expression using in situ hybridization (Fig. 6A(i)), and also detected *aadc* expression in the testes and vitellaria (Fig. 6A(ii), (iii)). Double FISH to detect *aadc* and somatic support cell markers in ovaries (*delta3, foxL,* and *ece1*) and testes (*ophis)* revealed that *aadc* is expressed in both female and male somatic gonadal cells (Fig. 6B, 6C). In ovaries, *aadc* expression was enriched in the *ece1^+^foxL^low^* cells at the periphery (Fig. 6B). To confirm this expression in somatic gonadal cells, we generated and validated polyclonal antibodies against AADC (Fig. S7C, S7D, and Methods). Immunofluorescence labeling with anti-AADC antibodies detected AADC within the somatic support cells, which form scaffolds surrounding the germ cells in ovaries and testes (Fig. 6D, 6E, Fig. S7D). These data suggest that AADC could act locally within somatic gonadal cells to produce monoamines. In support of this idea, we find differential expression of various serotonin synthetic enzymes and other pathway components in the ovary and testis transcriptomes (Fig. S7E).

**Figure 6.**
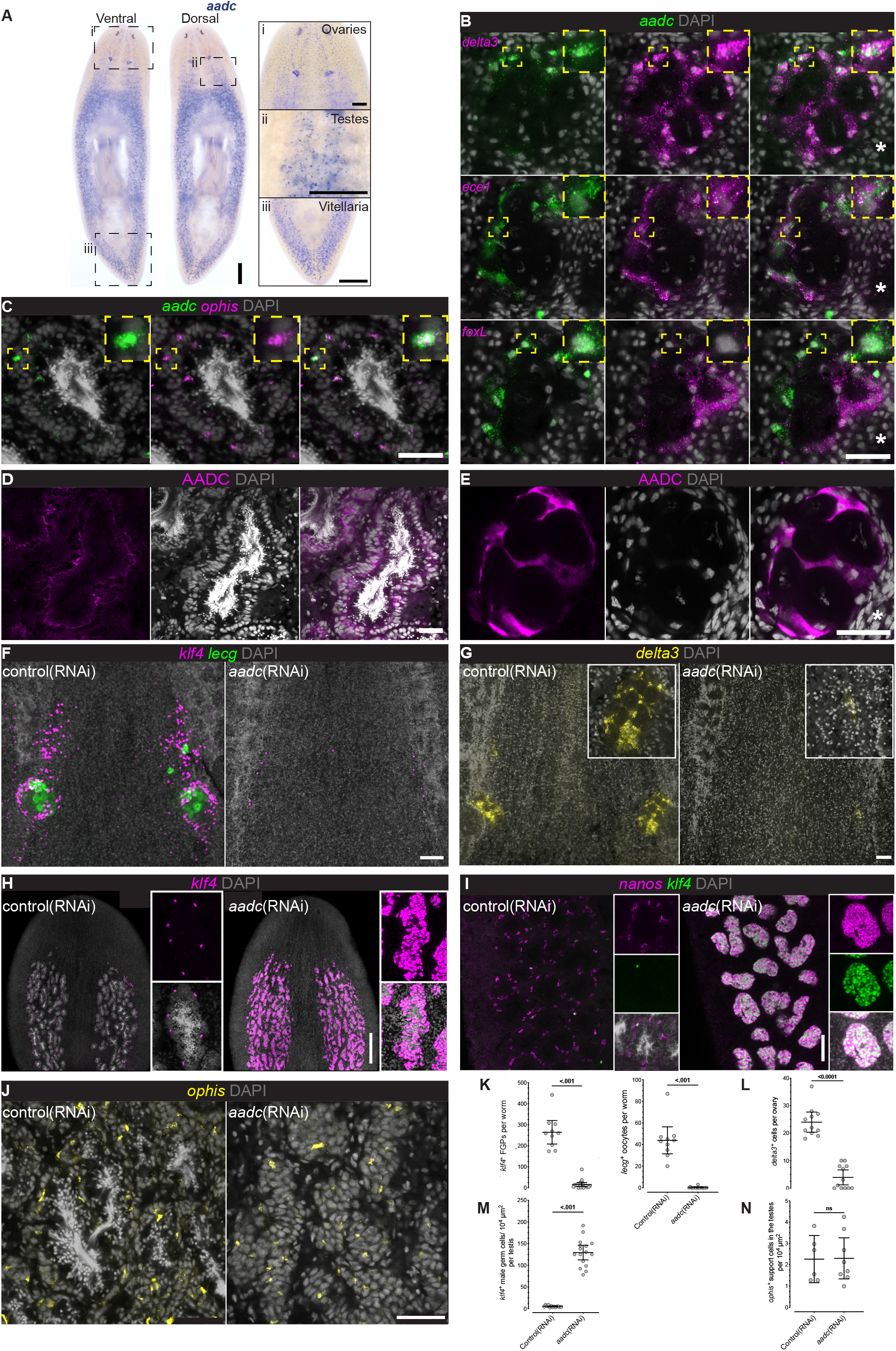
Opposing roles of AADC in female and male germ cell regeneration. (A) Expression of *aadc* (aromatic L-amino acid decarboxylase) is detected in the planarian reproductive system: (i) ovaries, (ii) testes, and (iii) vitellaria. (B) FISH to detect *aadc* and ovarian somatic markers (*delta3, ece1* and *foxL*) indicate *aadc* transcripts are enriched in the distal *ece1^high^foxL^low^* ovarian somatic cells. (C) FISH to detect *aadc* and testis somatic marker (*ophis)* indicates *aadc* expression in testis somatic support cells. (D, E) Immunostaining for AADC protein in testes (D) and ovaries (E). (F) FISH to detect *klf4^+^* FGPs and *lecg^+^* oocytes in control(RNAi) and *aadc*(RNAi) worms. (G) FISH for *delta3* to mark somatic ovarian cells in control and *aadc*(RNAi) worms. (H-I) FISH to detect *klf4* (H) and *nanos* (I), labeling early male germ cells in testes of control(RNAi) and *aadc*(RNAi) worms. (J) FISH for *ophis* to mark testis somatic support cells in control and *aadc*(RNAi) worms. (K-M) Quantification of *klf4^+^* FGPs and *lecg^+^* oocytes (K); *delta3^+^* ovarian support cells (L); *klf4^+^* early male germ cells (M); and *ophis^+^* male somatic cells (N), in control(RNAi) and *aadc*(RNAi) worms. Scale bars: (A,H) 500 μm; (B-E, J) 50 μm; (F, G, I) 100 μm. See also Figure S7.

To determine whether monoamines play a role in female germ cell development, we disrupted AADC function using the RNAi paradigm described above (Fig. 4A). Knockdown of *aadc* resulted in the failure to regenerate ovaries after amputation: we observed a dramatic reduction of *klf4^+^* FGPs and a complete absence of oocytes (Fig. 6F, 6K). Furthermore, *aadc*(*RNAi*) worms failed to regenerate the somatic compartment of the ovary (Fig. 6G, 6L). These results suggest that AADC activity is necessary for the specification of FGPs and implicates ovarian somatic cells in this process. More broadly, accessory reproductive organs (vitellaria, oviducts, sperm ducts, and the gonopore) failed to regenerate or be maintained, and egg-laying was abolished in *aadc*(RNAi) worms (Fig. S7F-H). No obvious *aadc* expression was detected in oviducts or sperm ducts (Fig. 6A), suggesting that AADC function within neurons, somatic gonadal cells, and/or vitellaria could act extrinsically in regeneration of these organs.

Surprisingly, *aadc* knockdown resulted in a novel testis phenotype. In contrast to the lack of ovaries after *aadc* knockdown, *aadc*(RNAi) animals had hyperplastic testes, consisting of early male germ cells (Fig. 6H). Typical testis organization is seen in control worms, with immature (*klf4^+^nanos^+^* and *klf4^−^nanos^+^*) germ cells located peripherally, and differentiating germ cells (spermatids and sperm) found towards the lumen of each lobe (Issigonis et al., 2021; Wang et al., 2007).The testes of *aadc*(RNAi) worms, however, were composed mainly of *klf4^+^nanos^+^* early male germ cells and lacked differentiated cells (Fig. 6H, 6I, 6M). Unlike the ovaries, somatic cells in the testes appeared unaffected after *aadc* knockdown (Fig. 6J, 6N). Taken together, these results implicate somatic gonadal cells as a potential source of monoamines, which may be acting differentially upon male and female germ cells in planarians.

## Discussion

Planarians exhibit extraordinary plasticity in reproductive development, including the ability to regenerate germ cells de novo; the mechanisms underlying this plasticity remain poorly understood. To uncover mechanisms underlying female germ cell development and regeneration, we generated gonad-specific transcriptomes. These studies identified genes defining progressive stages of female germ cell development and revealed heterogeneity of the somatic ovarian cells. In parallel, we discovered intriguing molecular similarities between female germ cells and neoblasts, the pluripotent stem cells that drive regeneration. We identified conserved, somatically expressed regulators, which have sex-specific functions in germ cell development and regeneration. Our findings underscore the key role played by somatic support cells not only in germ cell development but also in germ cell regeneration.

### Germ cell sex in a simultaneous hermaphrodite and the relationship of FGPs to pluripotent stem cells

Planarians are simultaneous hermaphrodites, developing ovaries and testes in different regions of the body. Because the genes implicated thus far in the earliest stages of germ cell development (*nanos* and *klf4*) are expressed in both male and female germ cells (Handberg-Thorsager and Saló, 2007; Issigonis et al., 2021; Sato et al., 2006; Wang et al., 2007), it was unclear whether the sex of these early germ cells was already determined. The identification of *zfs1* as a female-specific, germ cell-intrinsic factor indicates that planarian germ cells “know” their sex early in the course of their development. Presumably, inputs from global-patterning signals that define the animal’s major body axes (Reddien, 2018) are integrated such that germ cells born antero-ventrally toward the base of the brain adopt female fates, whereas those born dorso-laterally adopt male fates. Exploring the impact of global patterning signals on sex-specific germ cell specification will be an important future direction.

The distribution of FGPs in fields anterior to the ovaries, suggests two plausible scenarios. These fields could represent streams of migratory female germ cells that will later be incorporated into the ovary, similar to the example of planarian eye progenitors that are found in streams outside of the photoreceptors (Atabay et al., 2018; Lapan and Reddien, 2012). Alternatively, early female germ cells may be specified in a relatively broad antero-ventral region, but only those cells interacting with somatic gonadal cells (or other anatomical landmarks) will be incorporated into the ovary and produce oocytes. Whichever of these models best reflects reality, the gene expression changes we identified throughout female germ cell development suggest that FGPs/oogonia found around the margin of the ovary differentiate peripherally into oocytes that mature and move internally towards the tuba, where they exit from the ovary for fertilization (Fig. 7).

**Figure 7.**
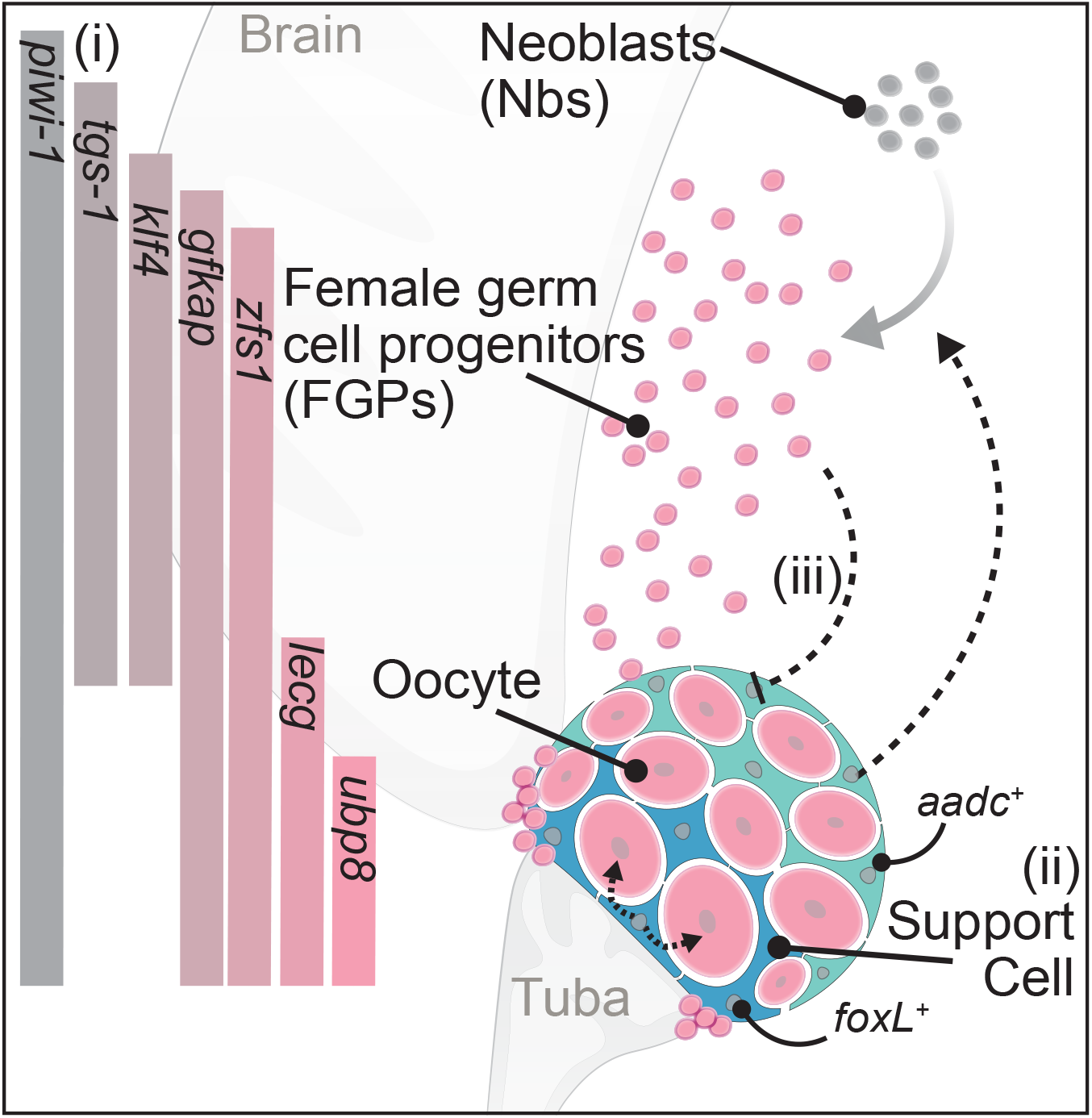
Schematic of planarian female germ cell differentiation and the roles of somatic gonadal cells. (i) Neoblasts specialize into FGPs antero-ventrally towards the base of the brain. FGPs around the margin of the ovary differentiate peripherally into oocytes that mature and move internally towards the tuba (fertilization duct). Gene expression at distinct stages of female germ cell differentiation is indicated on the left. Expression of a key female-specific factor, *zfs1*, indicates early and spatially restricted specification of germ cell sex in a simultaneous hermaphrodite. (ii) Somatic support cells are closely associated with germ cells within the ovary and display spatial heterogeneity in gene expression. Additionally, the tuba-distal *aadc^+^* somatic cells are required for regeneration of the female germ cells, whereas *foxL^+^* tuba-proximal cells regulate oocyte development, suggesting distinct functional domains within the ovarian somatic compartment. (iii) Loss of female germ cells triggers an expansion of ovarian somatic cells, suggesting that feedback from female germ cells regulates support cell numbers.

Previous studies have noted the similarities between neoblasts and germ cells, from the presence of ribonucleoprotein granules called chromatoid bodies, to the expression of several genes associated with germline development in other animals, including *vasa*, *piwi*, *tudor*, *pumilio*, and *bruno* (Guo et al., 2006; Reddien et al., 2005; Salvetti et al., 2005; Shibata et al., 1999; Solana et al., 2009). Single-cell transplantation into lethally irradiated hosts revealed a subpopulation of neoblasts capable of giving rise to all cell types in the animal; these pluripotent neoblasts are functionally defined as clonogenic neoblasts (cNeoblasts)(Wagner et al., 2011). Single-cell sequencing efforts to characterize a pluripotent neoblast subpopulation identified TSPAN-1 as a cell-surface marker that can be used to enrich for cNeoblasts (Zeng et al., 2018). These authors identified *tgs-1* and *pks1* (here renamed *gfkap*) as signature transcripts associated with this *tspan1*^+^ cluster in the asexual strain of *S. mediterranea*. We have shown here that in the sexual strain, these two genes are expressed abundantly in FGPs. Whereas *gfkap* is expressed at high levels constitutively in germ cells during homeostasis, it is only upregulated in neoblasts in response to wounding. This wound-responsive, neoblast-specific upregulation of genes associated with early stages of female germ cell development is particularly intriguing given the recent demonstration that neoblast fates are far more plastic than generally appreciated: specialized neoblast subclasses that express transcription factors associated with specific cell types are capable of generating daughters that adopt other fates (Raz et al., 2021). Our results suggest that this plasticity may involve transient activation of some germ cell-associated genes. Similarities in gene expression between germ cells and pluripotent neoblasts during regeneration are consistent with previous observations that planarian germ cells are capable of contributing to the regeneration of somatic tissues (Gremigni et al., 1980a, 1980b).

### Ovarian somatic support cells: key players in germ cell regeneration

Somatic support cells of planarian ovaries were observed ultrastructurally decades ago (Fischlschweiger, 1991, 1994; Gremigni and Falleni, 1998; Gremigni and Nigro, 1983; Harrath et al., 2011), but their functional characterization began with the discovery of *ophis,* a GPCR-encoding gene that is expressed in somatic gonadal cells and required for germ cell differentiation (Saberi et al., 2016). The paucity of these cells had thus far hindered their discovery by bulk and single-cell sequencing approaches (Fincher et al., 2018; Zayas et al., 2005). We used laser-capture microdissection to overcome this limitation, and the resulting gonad-specific transcriptomes enabled us to identify a set of sexually dimorphic ovarian support cell markers (*delta3, foxL,* and *ece1*). Their expression patterns revealed heterogeneity within ovarian somatic cells: enriched expression of *ece1* and *aadc* was observed in cells distal to the tuba, whereas enriched expression of *foxL* was detected in cells proximal to the tuba, associated with later-stage oocytes. We found that *aadc* was required for FGP specification and maintenance, whereas *foxL* was required for oocyte differentiation and maintenance, without affecting FGPs (Fig. 7). These disparate effects suggest that the tuba-distal and tuba-proximal somatic cell populations may control distinct stages of female germ cell development (Fig. 7). Such distinct functional domains would be consistent with the recently described functional compartmentalization of the somatic niche in the germarium of the *Drosophila* ovary (Shi et al., 2021; Tu et al., 2021) and the distinct populations of granulosa cells regulating reproductive onset vs duration in the mouse ovary (Niu and Spradling, 2020).

The expression patterns and knockdown phenotypes of the Notch signaling components reported here suggest that interactions between ovarian somatic cells (which express *delta3* and *notch4*) and accessory reproductive organs, the oviduct and tuba (which express *notch2* and *notch4*), help establish (and/or maintain) proper ovarian structure. Notch signaling also affects the germ cells: RNAi knockdown of *delta3* or *notch2* resulted in both the skewed distribution of FGPs toward the midline and an increase in the number of FGPs. The midline skewing could reflect the role of *delta3* in midline patterning: it is expressed at the midline and knockdown in the asexual strain leads to cyclopia (Sasidharan et al., 2017). We did not observe cyclopic worms in our RNAi regeneration assays, so it remains to be determined if FGPs are more sensitive to the disruption of midline cues than the photoreceptors, or if the altered distribution reflects altered signaling from the somatic ovary. Nonetheless, potential alterations in midline patterning seem unlikely to affect the number of FGPs; thus, the somatic ovary may be capable of communicating with FGPs and regulating their specification and/or proliferation at a distance.

*FoxL2* is a critical regulator of granulosa cell fate in mammals; homologs of this gene had yet to be characterized functionally in any invertebrate. Our observations that a planarian homolog (*foxL*) is expressed robustly in a subpopulation of somatic ovarian cells and is required for oocyte differentiation and maintenance suggest that the female-specific role of FoxL family members in the somatic gonad may well predate the emergence of vertebrates. The sexually dimorphic expression of planarian *foxL* is particularly intriguing in the context of this simultaneous hermaphrodite. In mice, female-specific FoxL2 and male-specific Dmrt1 transcription factors act in a mutually antagonistic manner to maintain gonadal sex: post-natal conditional knockout of *foxL2* results in granulosa cell transdifferentiation into Sertoli cells (Uhlenhaut et al., 2009); by contrast, post-natal conditional knockout of *dmrt1* results in Sertoli cell transdifferentiation into granulosa cells (Matson et al., 2011). In addition to *foxL*, planarians also have a *dmrt1* homolog (*dmd-1*), which is expressed specifically in male reproductive organs (including somatic cells of the testes) and is required for the specification, differentiation, and maintenance of male germ cells as well as accessory reproductive organs (Chong et al., 2013). Whether functional antagonism like that observed in mice, between female-specific *foxL* and male-specific *dmd-1* also plays a role in maintaining gonadal sex in this simultaneous hermaphrodite will be an important avenue for future studies.

### Potential non-neuronal roles of monoamines in germ cell development

Biogenic monoamines, like serotonin and dopamine, act as neurotransmitters across the animal kingdom (Weiger, 1997; Yamamoto and Vernier, 2011). It is perhaps less widely appreciated that many of these molecules predate the evolution of nervous systems (Roshchina, 2016) and play other, non-neuronal roles (Bellono et al., 2017; Lv and Liu, 2017; Matsuda et al., 2004). Our finding that AADC, an enzyme required for monoamine synthesis, plays sex-specific roles in planarian germ cell regeneration, implicates monoamines in the regulation of crucial stages of germ cell development, from specification to differentiation. AADC is involved in the synthesis of many different monoamines (including serotonin, tyramine, tryptamine, histamine, dopamine, and other catecholamines); thus, it is unclear whether the observed sex-specific effects reflect sex-specific activities of different monoamines or whether male and female tissues respond differentially to the same monoamine.

Although we do not yet know the identity of the monoamine(s) mediating the *aadc*(*RNAi*) phenotype, planarian gonads possess the machinery to respond differentially to the same monoamine. For example, we queried the gonad transcriptomes and found evidence for serotonergic pathways in both testes and ovaries (Fig. S7E). Tryptophan hydroxylase (TPH) catalyzes the conversion of L-tryptophan to 5-hydroxy-L-tryptophan, which is decarboxylated by AADC to produce serotonin. Vesicular monoamine transporters (VMAT) then package serotonin into vesicles for release, where it can bind serotonin receptors (5HTR). Serotonin transporters (SERT) terminate signaling by taking up extracellular serotonin. A homolog of the *Drosophila* peripheral tryptophan hydroxylase (TPH), *henna*, is highly expressed in both testes and ovaries, relative to the neuronally expressed *tph1* (Currie and Pearson, 2013; März et al., 2013; Sarkar et al., 2019). Planarian gonads also express *vmat* and *sert* homologs, suggesting the ability to release and recycle serotonin. Notably, ovaries and testes appear to differentially express different *vmats*, *serts*, and serotonin receptors. Thus, gonadal cells are capable of producing and responding locally to monoamines such as serotonin; the expression of distinct receptors in ovaries and testes suggests the possibility of differential responses. The expression of AADC by somatic support cells and the presence of specific monoamine receptors in both ovaries and testes supports the idea that these molecules can be produced and sensed locally within the gonads. Moreover, a recent report showed that serotonin could induce ovary development in another planarian species (*Dugesia ryukyuensis*), further supporting the role of monoamines in planarian reproductive system development (Sekii et al., 2019).

Might monoamines act in germ cell development in other animals? Genes encoding an entire serotonergic network, consisting of synthetic enzymes, receptors, and transporters are expressed in mammalian ovaries (Dubé and Amireault, 2007). In mice and humans, two paralogous genes encode Tryptophan Hydroxylase, the rate-limiting enzyme in serotonin synthesis. Expression of one paralog (*Tph1*) is found in a wide range of non-neuronal tissues and is responsible for synthesizing ∼95% of the body’s serotonin; whereas expression of the other (*Tph2*) is detected largely in neural cells (reviewed in Amireault et al. 2013). The embryonic defects observed in offspring of *Tph1*^−/−^ knockout females (Côté et al., 2007) are confounded by potential indirect effects from the compromised physiological states of these mice, which have diabetes and anemia (Amireault et al., 2011; Paulmann et al., 2009). These complications of interpreting *Tph1*^−/−^ phenotypes and the finding that the placenta serves as the source of serotonin that acts in the fetal forebrain (Bonnin et al., 2011) seem to have quelled investigations into potential roles of serotonin in oogenesis and early embryonic development. Furthermore, because *Tph2* is among the genes showing the greatest degree of upregulation in the female somatic gonad at the time of sex determination (Beverdam and Koopman 2006), it seems quite likely that it may act redundantly with *Tph1* in the ovaries. Similar expression of multiple serotonin receptors in the gonads would also likely compensate for loss of any single gonadally enriched gene. Thus, exploring potential roles of these deeply conserved molecules in the reproductive organs of a simpler animal, like the planarian, may overcome such concerns about redundancy and guide future tissue-specific knockout studies.

Regardless of whether or not the role of monoamines uncovered here reflects deeply conserved mechanisms for regulating germ cell development across animal phylogeny, the observed effects upon the planarian’s reproductive system have other important implications. Planarians are free-living representatives of the Phylum Platyhelminthes (the flatworms) and their reproductive system shares several important features with those of parasitic flatworms (flukes and tapeworms), which have major impacts on global public health (Collins and Newmark, 2013). All of these flatworms have ectolecithal eggs (i.e., yolk on the outside); yolk cells are produced by specialized accessory reproductive organs known as vitellaria and they are essential for embryonic development. Furthermore, transmission of parasitic flatworms requires prolific egg and yolk cell production. Because inhibition of *aadc* results in the loss of ovaries and other female accessory reproductive organs (including vitellaria), unraveling the role of monoamines in the female flatworm reproductive system may lead to new approaches for preventing parasite transmission.

## Acknowledgments

We thank Rosa Mejia Sanchez, John Brubacher, Melanie Issigonis, Jayhun Lee, Jiarong Gao, Tracy Chong, and Tania Rozario for critical manuscript feedback. We are grateful to past and present Newmark Lab members for valuable discussions. We especially thank David Forsthoefel (OMRF, Oklahoma) for sharing his initial work on optimizing LCM-RNAseq. We also thank Mayandi Sivagaru (Institute for Genomic Biology, University of Illinois at Urbana-Champaign [UIUC]) for LCM training, and Alvaro Hernandez (Roy J Carver Biotechnology Center, UIUC) for library preparation and Illumina sequencing. This work was supported by NIH Grant R01 HD043403. P.A.N. is an investigator of the Howard Hughes Medical Institute.

## Author Contributions

Conceptualization, Methodology, Writing, U.W.K and P.A.N; Investigation, Visualization, U.W.K; Funding Acquisition, P.A.N.

## Supplemental Figure Legends

**Figure S1.**
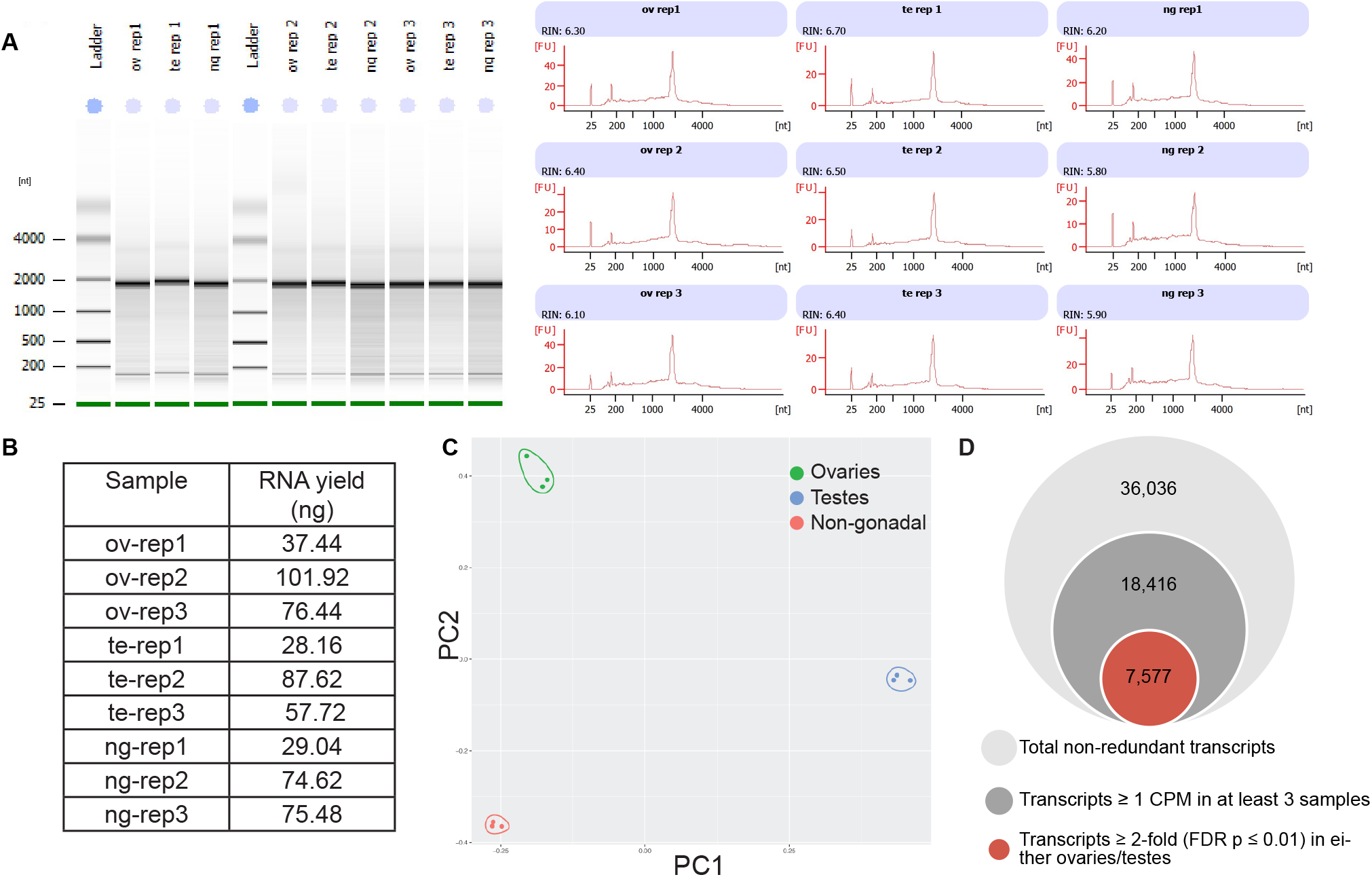
LCM-RNAseq approach to generate gonadal transcriptomes. (A) Bioanalyzer analysis of RNA integrity recovered from LCM-excised tissue. (B) RNA yields from LCM-dissected tissues (ov- ovary, te- testis, ng- non-gonadal, rep- replicate) (C) PCA analysis of LCM-RNAseq samples. (D) LCM-RNAseq results: 7,577 genes (≥ 2-fold; FDR p≤ 0.01) are preferentially expressed in ovaries or testes compared to non-gonadal tissue.

**Figure S2.**
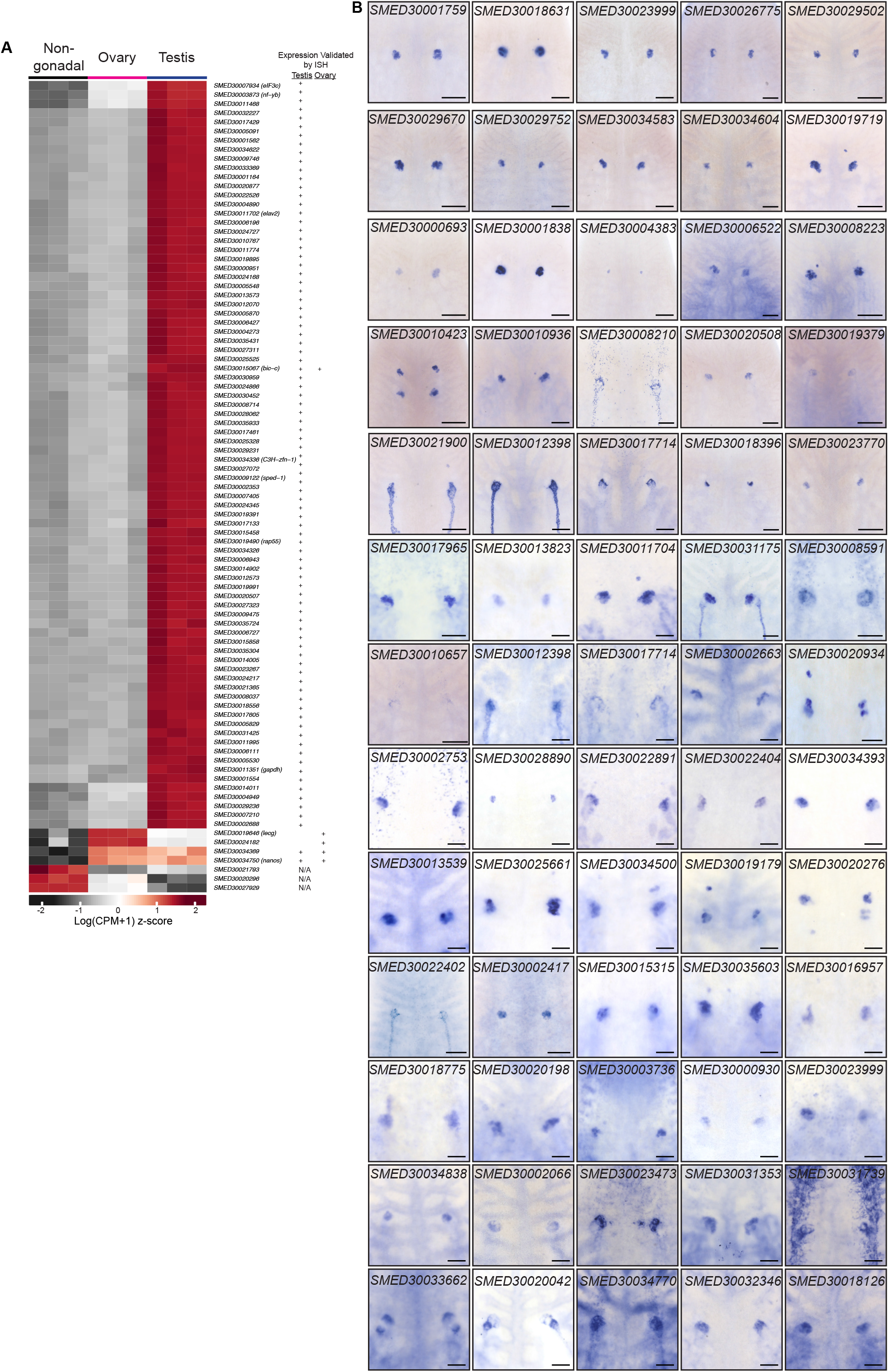
Validation of LCM-RNAseq generated gonadal transcriptomes. (A) Heatmap of 89 gonadal-enriched genes previously identified and validated (Wang et al. 2010). ISH results from Wang et. al study summarized in the table to the right. (B) Representative WISH for additional ovary-enriched candidate genes. Scale bars: 200 μm.

**Figure S3.**
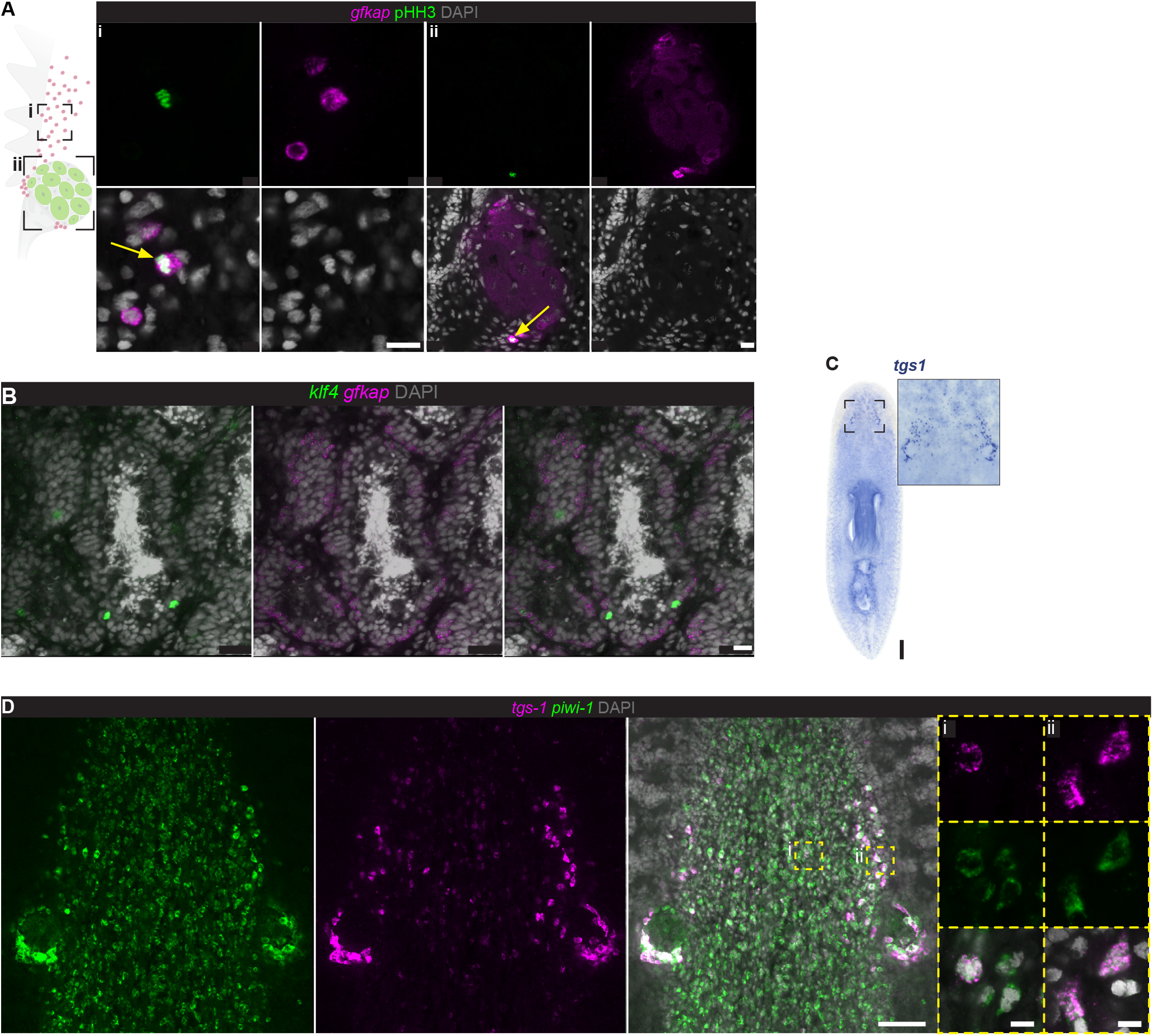
FGPs are a proliferative, extra-ovarian cell population, defined by coexpression of *klf4* and *gfkap*. (A) FISH to detect *gfkap* and anti-phospho histone H3 (pHH3) indicating proliferative activity of FGPs. (B) FISH to detect *gfkap* expression in testes. (C) WISH to detect *tgs-1*. (D) FISH to detect *tgs-1* and *piwi-1. Tgs-1* marks a pluripotent subset of neoblasts (i) and is enriched in FGPs (ii). Scale bars: (A,B) 20 μm; (C) 500 µm; (D) 100 µm; insets: 10 µm.

**Figure S4.**
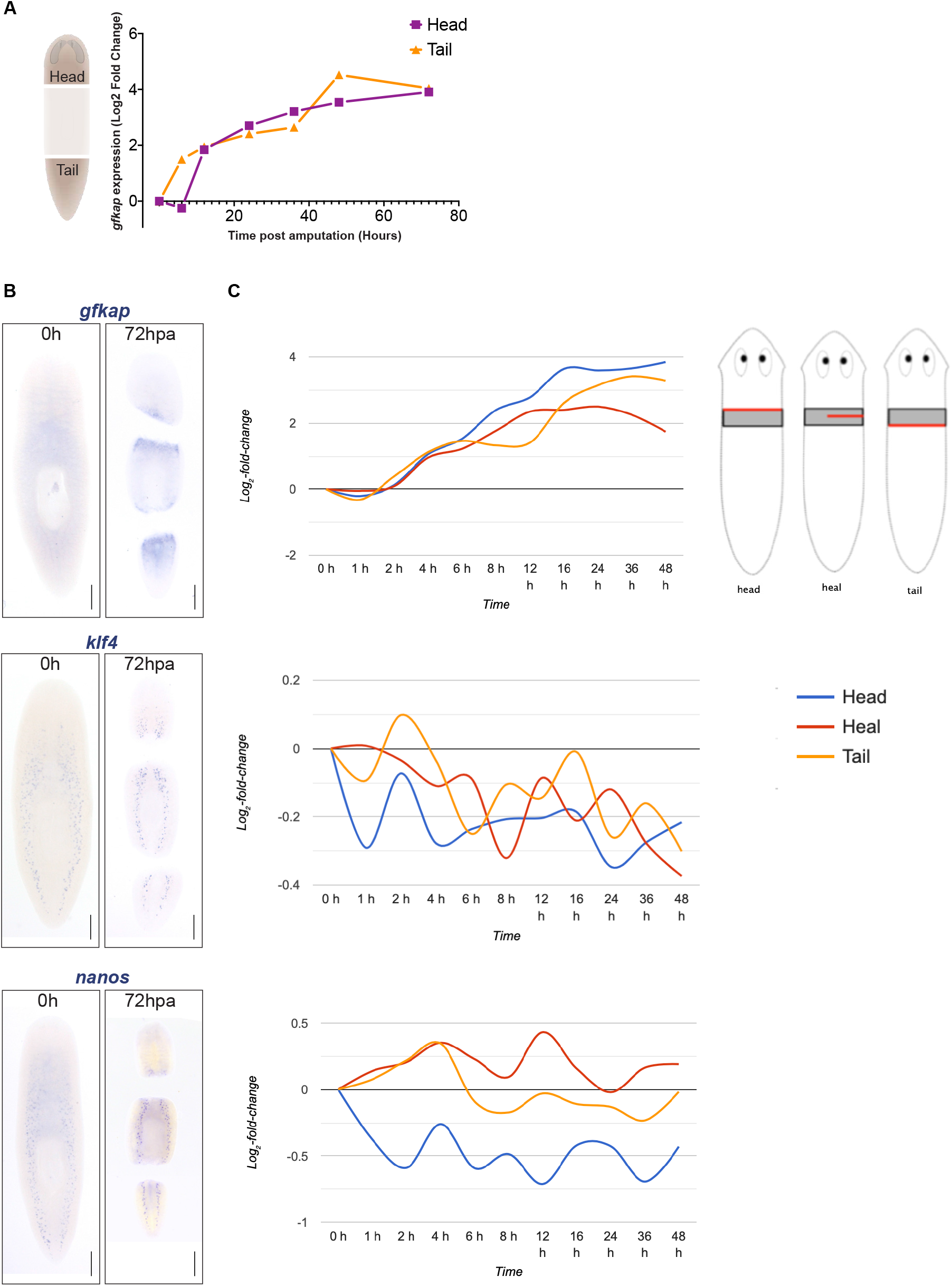
Expression of germ cell markers after wounding. (A) Expression of *gfkap* in head and tail fragments after amputation. Regeneration time course data (Kao et al., 2013) for *gfkap* retrieved from Planmine (http://planmine.mpi-cbg.de/)(Rozanski et al., 2019). (B) WISH for *gfkap, nanos* and *klf4* at 0 hours and 72 hours post-amputation (hpa) shows that only *gfkap* has detectable upregulation at wound sites after injury. (C) Expression of *gfkap, klf4* and nanos during a regeneration time course retrieved from Planmine (http://planmine.mpi-cbg.de/). Red: injury site; grey box: tissue analyzed.

**Figure S5.**
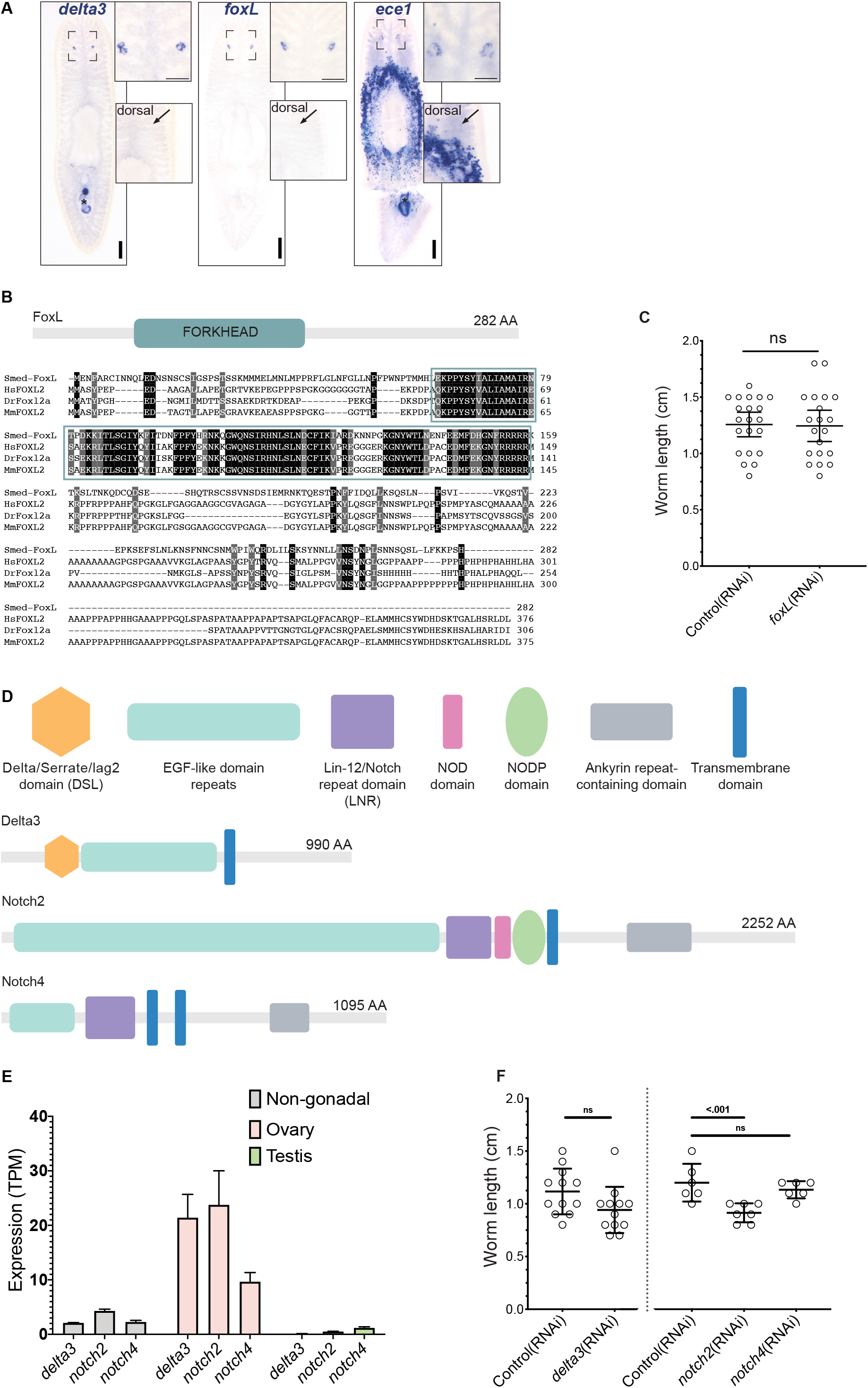
Analysis of genes expressed in ovarian support cells. (A) WISH for *delta3, foxL, and ece1* indicating expression in the ovary. Dorsal view shows that none of these genes have detectable expression in the testes (arrows pointing to the region where testes are found). ece1 and delta3 are detected in the penis papilla (asterisk), while *ece1* is highly expressed in the secretory cells around the pharynx as well. (B) Sequence alignment of Smed-FoxL with vertebrate FoxL2 proteins from Humans(Hs), Mouse(Mm) and Zebrafish(Dr). Black and gray shading indicate residues that are identical or conserved, respectively, among all four species. (C) Measurement of control(RNAi) and *foxL*(RNAi) worm length at the end of RNAi regeneration assay (ns: not significant; mean with 95% CI). (D) Domain architecture analysis of Smed-Delta3, -Notch2, and Notch4 using SMART and Interpro protein domain analysis tools. Smed-Delta3 contains a N-terminal Delta/Serrate/Lag2 (DSL) domain and EGF (epidermal growth factor) motifs. (E) *delta3*, *notch2*, and *notch4* mRNA expression levels (TPM-Transcripts per million) in the ovary, testis, and non-gonadal transcriptomes. (F) Measurement of control(RNAi), *delta3*(RNAi), *notch2*(RNAi), and *notch4*(RNAi) worm length at the end of RNAi regeneration assay (ns: not significant).

**Figure S6.**
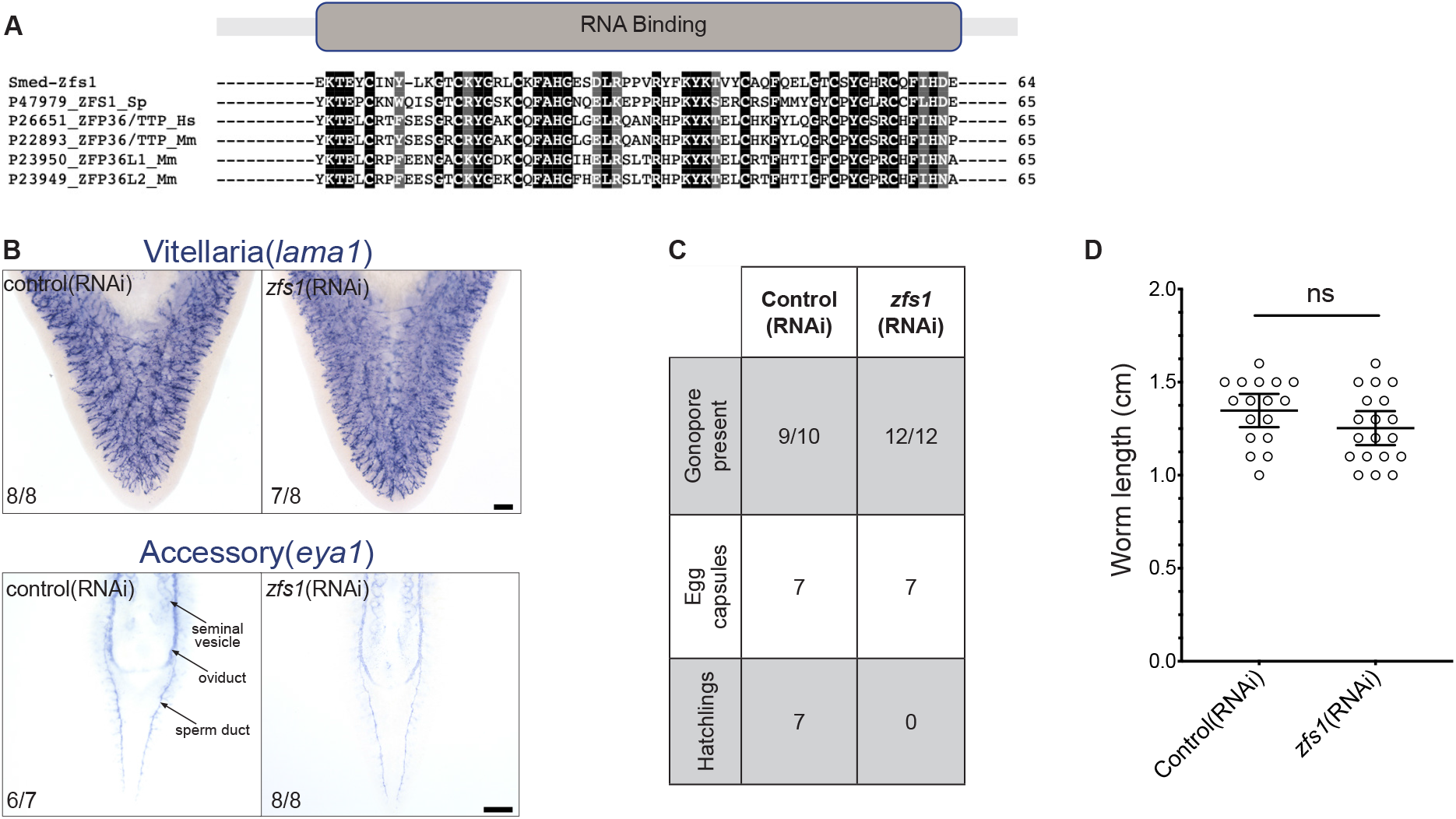
Female-specific role of *zfs1* in germ cell regeneration. (A) Sequence alignment of tandem zinc finger (C3H1) RNA binding domain of Smed-Zfs1 with homologs from Yeast (Sp), Humans(Hs) and Mouse(Mm). Black and gray shading indicate residues that are identical or conserved. (B) WISH for *lama1* (vitellaria), and *eya1* (accessory reproductive organs) in control and *zfs1*(RNAi) worms (tails regenerating a head). (C) Presence of gonopore, egg-laying, and egg-hatching observed for control and *zfs1*(RNAi) regenerated worms. (D) Measurement of control(RNAi) and *zfs1*(RNAi) worm lengths at the end of RNAi regeneration assay (ns: not significant). Scale bars: (B) 200 μm.

**Figure S7.**
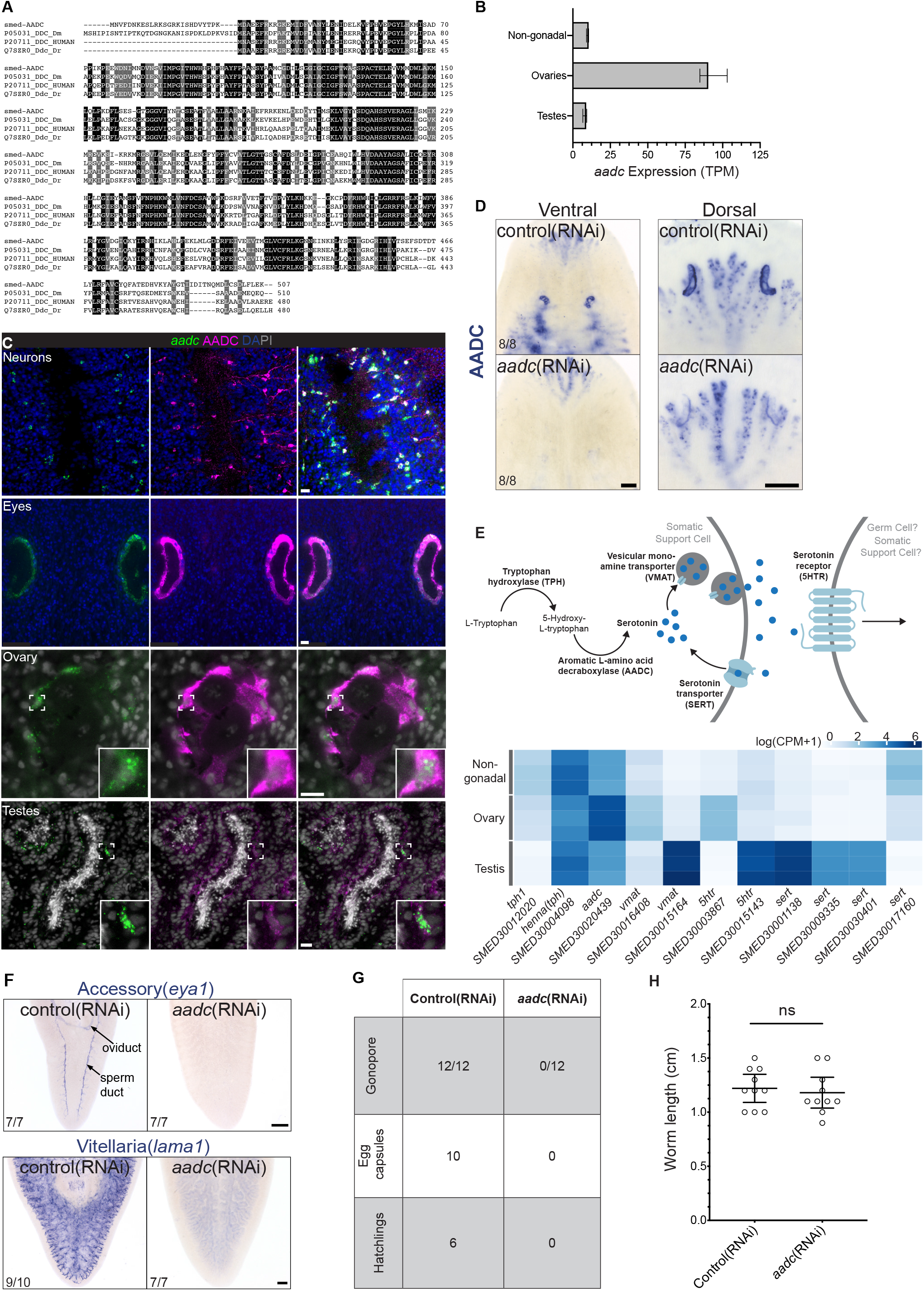
Roles of AADC in germ cell and reproductive development. (A) Sequence alignment of Smed-AADC with AADC proteins from Humans(Hs), Mouse(Mm) and Zebrafish(Dr). Black and gray shading indicate residues that are identical or conserved, respectively, among all four species.(B) *aadc* expression levels (TPM) in the ovary, testis, and non-gonadal transcriptomes. (C) Expression of AADC mRNA and protein in the indicated cell types and organs. (D) Labeling for AADC protein after *aadc* knockdown ablates labeling in the ovaries, eyes, and neurons, verifying the specificity of the AADC antibody staining in these organs. Non-specific labeling of gut branches can be seen in the background. (E) Schematic of canonical serotonin synthesis and signaling pathway in gonadal cells (Left). Normalized expression levels (LogCPM (counts per million)) of serotonin signaling pathway components in planarian gonads and non-gonadal tissues from the LCM-generated transcriptomes. (F) WISH for *lama1* (vitellaria), and *eya1* (accessory reproductive organs) in control and *aadc*(RNAi) regenerated worms (tails regenerating a head). (G) Presence of gonopore, egg-laying, and egg-hatching observed for control and *aadc*(RNAi) regenerated worms (tails regenerating a head). (H) Measurement of control(RNAi) and *aadc*(RNAi) worm lengths at the end of RNAi regeneration assay (ns: not significant). Scale bars: (C) 20 μm; (D, F) 200 μm.

## Methods

### Planarian Maintenance and Care

Sexual *S*. *mediterranea* were maintained at 18°C in 0.75X Montjuïc salts (Cebrià and Newmark, 2005) supplemented with 50 ug/mL Gentamicin (Gemini Bio-Products) and fed beef liver paste. Animals were stored in Ziploc containers or Petri dishes for RNAi. Animals were starved at least one week before any experiments.

### Laser-capture Microdissection and RNA extraction

Planarians were killed in chilled 2% HCl in PBS (RNAse-Free) and then fixed in 100% Acetone for 1 hour at −20°C. Planarians were then incubated in 10, 20, 30% sucrose in PBS (RNAse-Free) for 20 minutes each before embedding in Tissue Freezing Medium (TFM) blocks (Ted Pella). The samples were then cryosectioned at 16 µm thickness onto PEN membrane slides (Thermo Fisher). The slides were stained with 1% cresyl violet and 1% eosin Y (Sigma) in 75% ethanol (Clément-Ziza et al., 2008). The tissue samples were dissected using Veritas Laser Capture Microdissection System (Arcturus) and captured on to Capsure HS LCM Caps (Thermo Fisher). After capture, RNA was isolated using the Arcturus Picopure RNA isolation kit (Thermo Fisher). Ovarian tissue samples were collected from 14 worms (10 worms used for cross-sections and 4 worms used for sagittal sections) for each replicate (3 replicates total). Testis and non-gonadal tissue samples were collected from 6 worms for each replicate (4 worms used for cross-sections and 2 worms used for sagittal sections). RNA concentrations were determined using Qubit Fluorometer (Thermo Fisher), and RNA integrity was analyzed using 2100 Bioanalyzer Instrument (Agilent).

### RNA sequencing and gene expression analysis

Libraries were generated using Trueseq RNA stranded kit (Illumina), and 100 nt single-end sequencing reads were generated on HiSeq 2500 (Illumina). Trimming of adapters and low-quality reads as well as subsequent read mapping and differential gene expression analyses were performed using CLC Genomics Workbench (Qiagen). Reads were mapped to a de novo assembled *S. mediterranea* transcriptome (Smed2014) (Robb et al., 2015) with the *Smed-zfs1* transcript added. The transcriptome was annotated in OmicsBox (BioBam) using the SwissProt Protein database. Heatmaps were generated in R using the ComplexHeatmaps package (Gu et al., 2016).

### Molecular Biology Methods

200–1000 bp fragments of the gene of interest (Table S2) were amplified using Platinum Taq DNA Polymerase (Invitrogen) from cDNA. Amplified fragments were cloned into pJC53.2 (Collins et al., 2010) via TA cloning. Riboprobes and dsRNA were synthesized as previously described (King and Newmark, 2013; Rouhana et al., 2013).

### In situ hybridization and Immunohistochemistry

Whole-mount ISH was performed as previously described (King and Newmark, 2013) with modifications for larger sexual worms: formaldehyde fixation was increased to 30 min, proteinase K treatment was increased to 20-25 min, and post-proteinase K fixation was increased to 30 min. pHH3 was labeled using anti-phospho-Histone H3 (Ser10) (Millipore Sigma). AADC polyclonal antibody was generated by injecting a synthetic peptide (DVYTPKMDAEEFRKRGKE) into rabbits (Pierce Biotechnology, Rockford, IL). The serum was affinity-purified with the peptide antigen and used at a dilution of 1:2000 for immunohistochemistry. Colorimetric and FISH/immunofluorescence samples were imaged on Axio Zoom.V16 (Carl Zeiss) and LSM 710 or 880 confocal microscope (Carl Zeiss), respectively. Cell counts were performed using the spot detection tool in Imaris (Bitplane).

### RNA interference

Knockdowns were performed by feeding in vitro-transcribed dsRNA, as previously described (Rouhana et al., 2013). 6-8 mature sexual animals were fed ∼10-20 ug dsRNA mixed with 90 uL liver puree: water mixture 5:1 (with food coloring) in a petri dish. For all RNAi experiments, dsRNA corresponding to bacterial *ccdB* gene was used for negative controls. Animals were fed dsRNA 4 times, cut posterior to the ovaries, and tail fragments were allowed to regenerate for two weeks, followed by 6-8 dsRNA feedings.

### Protein domain and Phylogenetic analysis

Conserved protein domains were analyzed using InterProScan, SMART and Phobius protein domain analysis tools. Predicted protein sequences were aligned using MUSCLE.

### Statistical analysis

All two-sample and three sample comparisons were analyzed using Welch’s t-test or one-way ANOVA (Dunnett’s test), respectively, in Prism (GraphPad). Differential expression for RNAseq was analyzed in CLC Genomics Workbench (Qiagen) using the Wald test.

